# A massively parallel reporter assay reveals focused and broadly encoded RNA localization signals in neurons

**DOI:** 10.1101/2021.04.27.441590

**Authors:** Martin Mikl, Davide Eletto, Minkyoung Lee, Atefeh Lafzi, Farah Mhamedi, Simona Baghai Sain, Kristina Handler, Andreas E. Moor

## Abstract

Asymmetric subcellular localization of mRNA is a common cellular phenomenon that is thought to contribute to spatial gene regulation. In highly polar neurons, subcellular transcript localization and translation are thought to enhance cellular efficiency and timely responses to external cues. Although mRNA localization has been observed in many tissues and numerous examples of the functional importance of this process exist, we still lack a systematic understanding of how the transcript sorting machinery works in a sequence-specific manner.

Here, we addressed these gaps by combining subcellular transcriptomics and rationally designed sequence libraries. We developed a massively parallel reporter assay (MPRA) for mRNA localization and tested ~50,000 sequences for their ability to drive RNA localization to neurites of neuronal cell lines. By scanning the 3’UTR of >300 genes we identified many previously unknown localization regions and mapped the localization potential of endogenous sequences. Our data suggest two ways the localization potential can be encoded in the 3’UTR: focused localization motifs and broadly encoded localization potential based on small contributions.

We identified sequence motifs enriched in dendritically localized transcripts and tested the potential of these motifs to affect the localization behavior of an mRNA. This assay revealed sequence elements with the ability to bias localization towards neurite as well as soma. Depletion of RNA binding proteins predicted or experimentally shown to bind these motifs abolished the effect on localization, suggesting that these motifs act by recruiting specific RNA-binding proteins.

Based on our dataset we developed machine learning models that accurately predict the localization behavior of novel sequences. Testing this predictor on native mRNA sequencing data showed good agreement between predicted and observed localization potential, suggesting that the rules uncovered by our MPRA also apply to the localization of native transcripts.

Applying similar systematic high-throughput approaches to other cell types will open the door for a comparative perspective on RNA localization across tissues and reveal the commonalities and differences of this crucial regulatory mechanism.

## Introduction

The cytoplasm is a tightly regulated space that accommodates countless parallel tasks in specialized compartments. Asymmetric subcellular mRNA distributions have been observed in a variety of polar cell types (Das et al., 2021; Lécuyer et al., 2007; Mili et al., 2008; Moor et al., 2017). mRNA localization might be an energy-efficient way to generate corresponding protein gradients, it might prevent harmful protein effects by ectopic activity or accumulation, and it could accelerate cellular response to extrinsic stimuli by activation of localized protein translation (Blower, 2013).

Neurons show a high degree of functional compartmentalization, which to a large part is achieved through transporting specific transcripts into dendrites or axons, where they are available for local (and sometimes activity-dependent) translation (Glock et al., 2017). RNA localization in neurons was first demonstrated using in situ hybridization techniques (Burgin et al., 1990; Kleiman et al., 1990) and also revealed differences in the localization characteristics between neuronal cell types and brain regions (Paradies and Steward, 1997). Fractionation-based approaches allowed for the characterization of the entire pool of dendritically localized mRNAs (Gumy et al., 2011; Taylor et al., 2009), revealing the richness of the local transcriptome and subsequently proteome (Cajigas et al., 2012; Zappulo et al., 2017) as well as isoform-specific regulation (Ciolli Mattioli et al., 2019). Recent advances in spatial transcriptomics have enabled the study of dendritically localized RNAs at single cell resolution (Perez et al., 2021), with massively parallel hybridization approaches (Wang et al., 2020), and with expansion sequencing (Alon et al., 2021). This body of work reveals a wealth of mRNA localization patterns. The extent of this phenomenon in steady-state and its dynamic adaptation in physiology (Jambor et al., 2015) suggest that RNA localization is a tightly regulated process.

Most studied RNA localization phenotypes depend on interactions between RNA binding proteins (RBP) and the 3’ untranslated region (3’UTR) of a gene (Buxbaum et al., 2015). Approximately 1500 RBPs, interacting with a variety of RNA molecules, have been identified in mammalian cells (Van Nostrand et al., 2020). Several well-characterized localized transcripts are linked by RBPs to molecular motors, which enable active transport on the cytoskeleton (Holt and Bullock, 2009). Alternatively, localized RBPs have been shown to prevent RNA degradation (Medioni et al., 2012) or to capture and anchor transcripts and thereby create subcellular asymmetry in RNA localization (Buxbaum et al., 2015). The localization of beta-actin mRNA to the leading edge in fibroblasts and to axonal growth cones in neurons are well-studied examples of RNA localization (Bassell et al., 1998): they have been found to be mediated by the RBP ZBP1 binding to a “zip-code” in the three prime untranslated region (3’UTR) of the beta-actin mRNA (Patel et al., 2012). Apart from a very limited number of such cases with a transparent link between a specific motif and a trans-acting factor mediating RNA sorting, the link between sequence and localization potential remains obscure and we still lack a systematic understanding of how the transcript sorting machinery works in a sequence-specific manner.

Large scale testing of rationally designed or random sequence libraries has immensely contributed to elucidating the regulatory grammar of transcription (Arnold et al., 2013; Grossman et al., 2017; Kheradpour et al., 2013; Patwardhan et al., 2012; Sharon et al., 2012), splicing (Adamson et al., 2018; Mikl et al., 2019; Rosenberg et al., 2015; Soemedi et al., 2017; Wong et al., 2018), polyadenylation (Bogard et al., 2019; Vainberg Slutskin et al., 2019), miRNA-mediated regulation (Slutskin et al., 2018), other forms of translational control (Mikl et al., 2020; Weingarten-Gabbay et al., 2016), and RNA nuclear enrichment and export (Lubelsky and Ulitsky, 2018; Shukla et al., 2018; Zuckerman et al., 2020) demonstrating the power and universal applicability of such approaches. Although they have become an experimental pillar of studies in gene regulation, a dedicated high-throughput systematic attempt to dissect the regulatory logic of subcellular RNA localization is still lacking.

Here, we address our gaps in understanding the regulatory code of mRNA localization in neurons by combining subcellular transcriptomics and massively parallel reporter assays. This enabled us to functionally test endogenous and synthetic localization elements with unprecedented scale, determine how the localization potential is encoded in the 3’UTR sequence, identify protein mediators of localization, and train computational models to predict the localization of novel sequences.

## Results

### A massively parallel reporter assay for RNA localization in neurons

To dissect the sequence-encoded regulation of RNA localization in a systematic manner we developed a reporter system that would allow us to test tens of thousands of potential regulatory sequences for their ability to drive localization to neurites.

To select candidate sequences to test, we built on earlier studies characterizing the neurite- and soma-enriched transcriptome (Ciolli Mattioli et al., 2019; Middleton et al., 2019; Taliaferro et al., 2016; Zappulo et al., 2017). In general, the overlap in dendritically localizing RNAs reported in these studies is very low (Middleton et al., 2019), which could be due to differences in the experimental system or cell type used. We selected a core set of dendritically localizing RNAs identified in several independent studies (Middleton et al., 2019) and added genes that were found to be enriched in at least one of the published neurite transcriptome datasets (Methods). In addition, we selected 5 RNAs which have consistently been reported as soma-restricted in previous studies (Rragb, St6gal1, Gpr17, Ogt, Pgap1). This resulted in a set of 324 3’UTRs which formed the basis for designing a synthetic oligonucleotide library of altogether 47,989 sequences (Table S1).

Oligonucleotides in our library comprised common primers for amplification, a unique barcode and a 150 nt long variable region containing a 3’UTR sequence (Figure 1A). They were synthesized (Twist Bioscience), PCR amplified, and cloned downstream of a reporter gene (GFP). We transfected the final library into two mouse neuronal cell lines (CAD and Neuro-2a cells), which were differentiated into a neuron-like state by serum starvation (see methods, Figure S1). To allow physical separation of soma and neurites, cells were grown on microporous membranes coated with matrigel on the bottom, such that neurites would extend into the bottom compartment while cell bodies remained on top of the membrane. This experimental system has been previously used for sequencing the native soma and neurite transcriptome in CAD and Neuro-2a cells (Taliaferro et al., 2016) as well as in mESCs differentiated into neurons (Zappulo et al., 2017).

**Figure 1.**
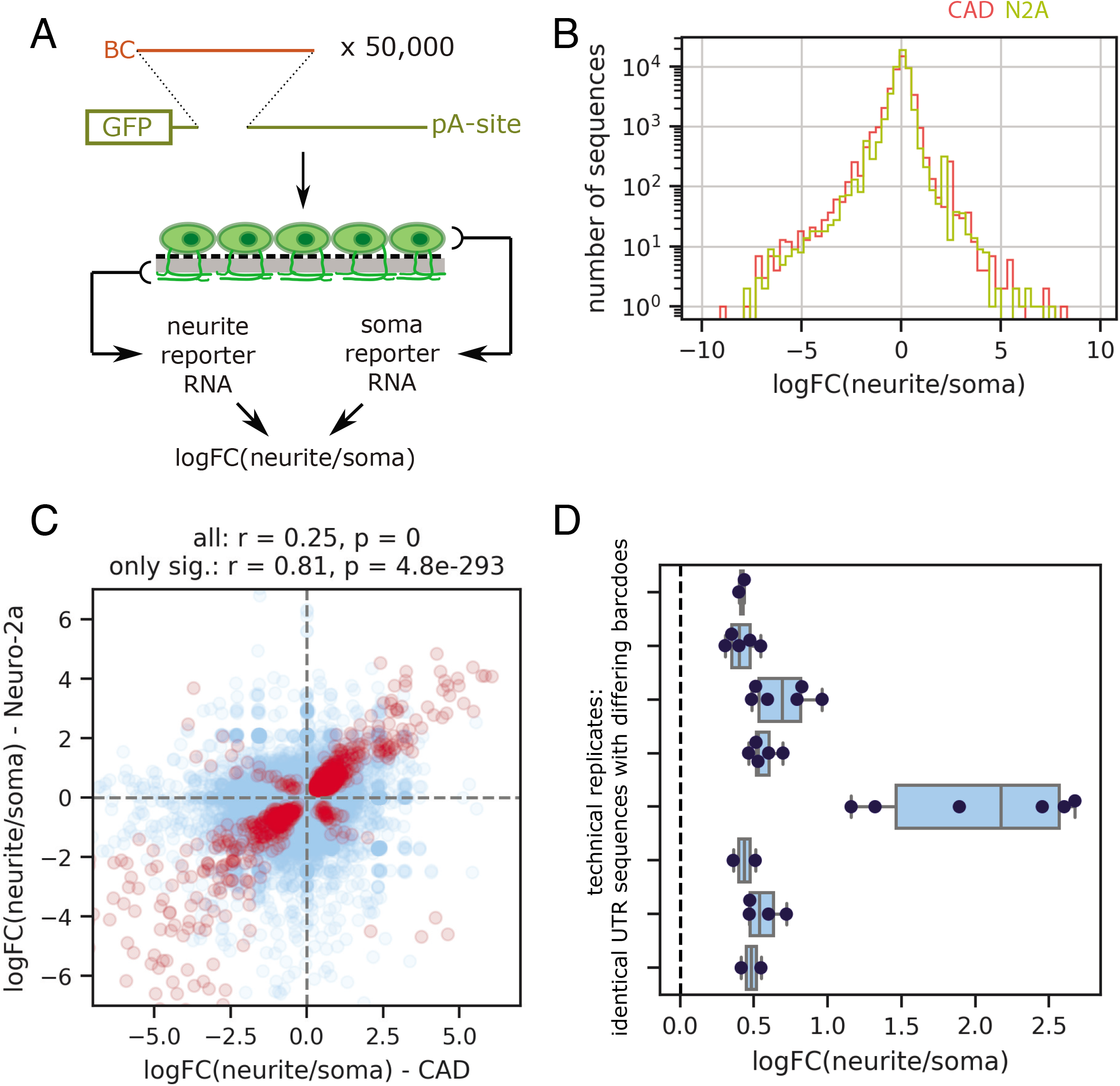
A massively parallel reporter assay for RNA localization in neurons. A. Experimental outline. An oligonucleotide library (containing a 5’ barcode (12 nt) and a 150 nt variable 3’UTR region is cloned downstream of the *gfp* coding sequence and upstream of a polyadenylation site. The final reporter library is transfected into neuronal cells, and reporter RNA from soma and neurite compartments are collected and sequenced, resulting in a measure for the enrichment of each library variant in the neurite vs. soma compartment; BC: barcode; pA-site: polyadenylation site. B. Density plot of logFC(neurite/soma) measured in CAD (red) and Neuro-2a cells (yellow); n=47347 for both. C. logFC(neurite/soma) measured in CAD cells are plotted against the logFC measured for the same sequence in Neuro-2a cells, for all sequences tested (light blue, n=47347) or only those with statistically significant enrichment (neurite or soma, p=0.05) in both cell lines (red, n=3412). D. logFC(neurite/soma) for groups with identical native 3’UTR sequence, but different barcodes; pooled measurements for CAD and Neuro-2a cells.

Total RNA from soma and neurite fractions was extracted from three biological replicates and cDNA was synthesized, including incorporation of unique molecular identifiers (UMI). Reporter library sequences were amplified by PCR and subjected to Illumina sequencing to determine the relative enrichment (log2 fold-change (logFC)) of each library sequence in the neurite and soma fraction, respectively (Methods).

In both cell lines most sequences did not show a preferential enrichment in the soma or neurite fraction (Figure 1B). Without active retention in the soma, RNAs can diffuse into (proximal) dendrites; therefore we expect to find most RNAs in both fractions. Significant enrichment of a sequence in the neurite fraction, however, is indicative of active transport or local anchoring or protection from degradation in neurites. In line with the stochastic nature of the distribution of RNAs without active transport, the general correlation between the two cell lines was modest (Pearson r=0.25, p=0, Figure 1C, light blue). However, when comparing only sequences with significant enrichment in either neurite or soma, the concordance between the two cell lines was much higher (Pearson r=0.81, p=4.8×10^-293^, Figure 1C, red), suggesting that the same sequences are actively sorted into neurites (positive log fold-change) or retained in the soma (negative log fold-change). Comparing logFC measured in our assay for library variants sharing the same sequence and differing only in the barcode showed good concordance in the direction of the enrichment (soma or neurite) and its magnitude (logFC, Figure 1D).

### 3’UTR fragments recapitulate the localization behavior of the endogenous transcript

In order to map sequences driving localization, we scanned the 3’UTR of 311 genes resulting from our analysis of published datasets (see above). For each gene, our oligonucleotide library contained tiles of length 150 nucleotides (nt), covering the entire 3’UTR with a step size (i.e. distance between start positions of adjacent tiles) of 50 nt. In our MPRA we tested the resulting 13,753 3’UTR tiles both in CAD and Neuro-2a cells, providing a landscape of localization potential along the 3’UTR (Figure 2A). Based on the localization behavior of these 150 nt tiles, we could in some cases identify a segment of the 3’UTR that showed strong enrichment in the neurite fraction and could constitute the neurite-targeting element mediating localization of the entire native 3’UTR, e.g. in Shank1, Camk2a, Vapb and others (Figure 2B,C, Figure S2). Prior work has analyzed the dendrite-localizing potential in the Camk2a gene and annotated most of it to a sequence stretch at the end of its 3’UTR that is only present in the longest Camk2a isoform (Tushev et al., 2018). Our Camk2a tile measurements (Figure 2C) validate these prior findings and resolve the localization potential from a previous 1kb resolution down to a 200bp stretch at the very end of the 3’UTR.

**Figure 2.**
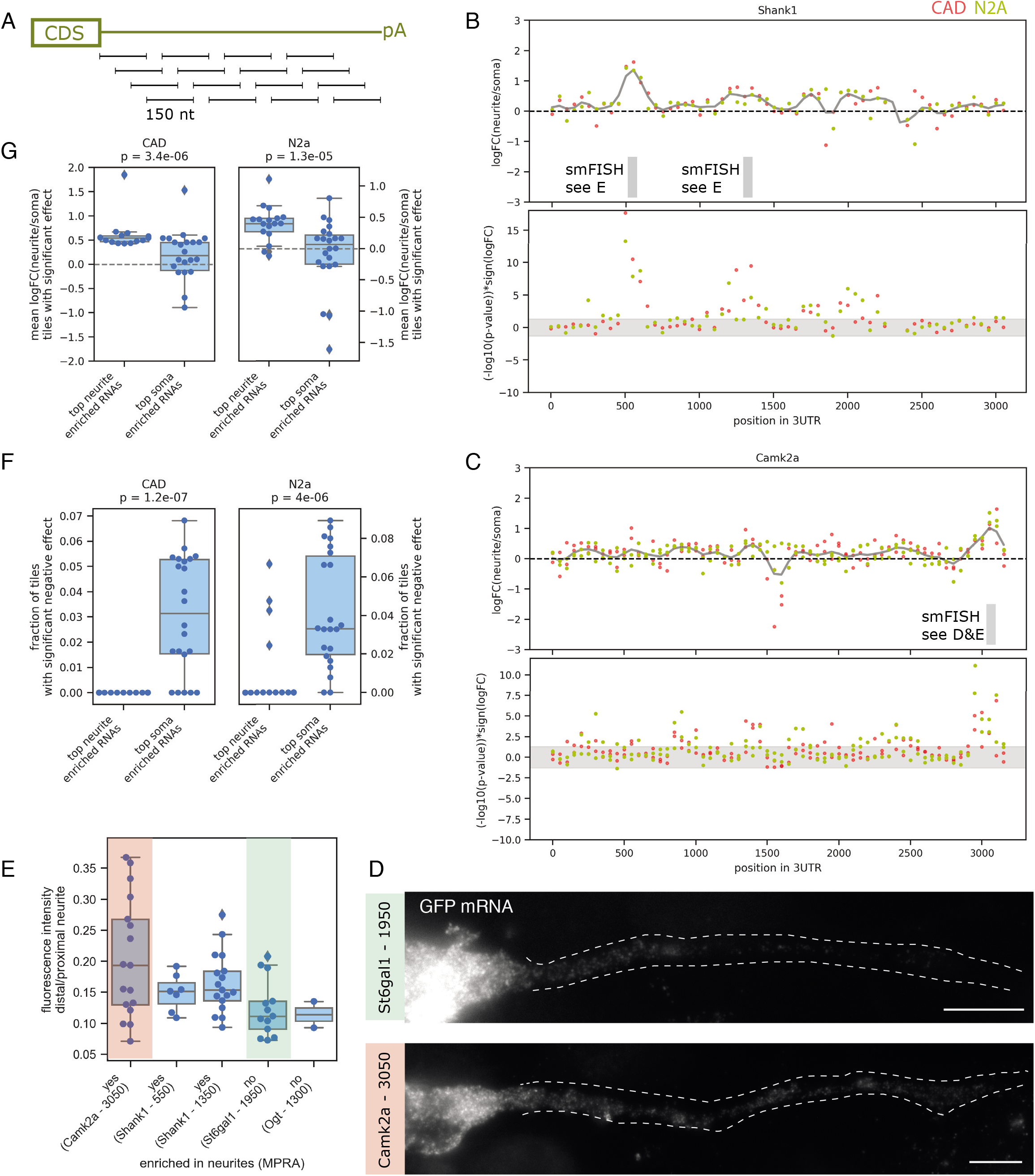
3’UTR fragments recapitulate the localization behavior of the endogenous transcript. A. Tiling of the 3’UTR of endogenous genes; starting immediately after the stop codon, segments of 150 nt are tested in isolation, with a distance of 50 nt between two adjacent tiles thereby creating a sequence overlap of 100 nt between adjacent tiles. BC. Measurements (upper panel: logFC(neurite/soma), lower panel: -log_10_(p-value) of the effect, multiplied by the sign of the logFC) for tiles along the 3’UTR of Shank1 (B) and Camk2a (C), as measured in CAD (red) and Neuro-2a (yellow) cells; the gray horizontal band denotes the area with p>0.05; the positions for which smFISH validations are shown in panels D and E are indicated below. DE. smFISH images (D) and quantification (E) for selected library sequences, the imaged and quantified individual 3’UTR reporters correspond to the tiles indicated in panels B and C. F. Each data point represents the mean logFC(neurite/soma) of all tiles corresponding to a segment taken from the same native 3’UTR and showing statistically significant enrichment in any compartment (neurite or soma, p=0.05); the groups correspond to the overlap of the 30 genes with the highest neurite (“top neurite enriched RNAs”) or soma enrichment (“top soma enriched RNAs”) in both CAD and Neuro-2a cells as measured by Taliaferro et al. G. Each data point represents the fraction of all tiles corresponding to a segment taken from the same native 3’UTR that showed statistically significant enrichment in the soma compartment; the groups correspond the overlap of the 30 genes with the highest neurite (“top neurite enriched RNAs”) or soma enrichment (“top soma enriched RNAs”) in both CAD and Neuro-2a cells as measured by Taliaferro et al.

To validate the results of our MPRA we constructed individual reporters carrying either soma restricted or neurite localizing 3’UTR tiles. We transfected these reporters into differentiated CAD cells and performed single molecule fluorescence *in situ* hybridization (smFISH) using probes targeting the *gfp* coding sequence (Methods). The localization behavior observed for individual library sequences generally matched the results from the MPRA, in particular when comparing distal portions of neurites which are less likely to be affected by passive diffusion of the RNA due to high expression levels (Figure 2D,E).

While the 3’UTRs described above indicated that the localization potential is encoded by a defined region, in other cases the localization potential was less clearly localized; instead, many 3 ‘UTR regions showed a moderate tendency for neurite enrichment, such as in the case of Shank3, Cntn2 and others (Figure S3). We hypothesize that the localization potential in these, or potentially in most, RNAs is broadly encoded, with many small contributions making up the net localization behavior of the native 3’UTR. To corroborate this hypothesis we compared the localization behavior of the tiles to the localization of the entire native 3’UTR as measured using a similar experimental setup (Taliaferro et al., 2016). Specifically, we selected sets of robustly neurite- vs. soma-enriched genes and compared the MPRA-based neurite/soma enrichment of the tiles mapping to these genes. Interestingly, the mean of the logFC of all tiles mapping to a gene was indicative of the localization behavior of the corresponding native transcript (Figure S4A). The separation between neurite and soma enriched genes further increased when only taking into account tiles with a significant enrichment in either the neurite or the soma fraction (Figure 2F).

We then analyzed the localization behavior of the neurite-enriched tiles mapping to a specific gene, without taking into account whether they map to one defined localization region or whether they are spread out over the transcript. This showed no visible difference between genes with a distinct localization element (Figure S4B, left) and genes with broadly encoded localization potential (Figure S4B, middle). Both groups exhibited a similar tendency for significant positive logFCs of their 3’UTR tiles, in contrast to soma-localized genes (Figure S4B, right), which did not show a tendency for more neurite-enriched tiles. Interestingly, a soma-enriched tile was a strong predictor of the localization behavior, as most top localizing 3’UTRs did not contain a tile exhibiting soma enrichment (Figure 2G, Figure S4A,B).

In contrast, soma-restricted RNAs showed less significant enrichment at the level of individual tiles, and if a gene contained a tile with enrichment in neurites then it also contained one enriched in the soma fraction, leading to no net enrichment (Figure S5). Taken together, these data indicate that the localization behavior of a transcript is determined by integrating over many effects, potentially with small sizes.

### Single RBP motifs can only prevent, but not enforce neurite localization

In order to dissect the mechanisms and trans-acting factors that mediate RNA localization to dendrites, we created an additional library of 34,236 sequences, which was cloned and tested in the same experimental pipeline as above. Here, we used 5841 150 nt regions taken from the 3’UTR of 230 genes to test the potential of RBP binding sites to drive localization of the transcript.

We analyzed existing neurite and soma transcriptomic data (Middleton et al., 2019; Zappulo et al., 2017) to identify RBP motifs preferentially found in neurite- or soma-enriched transcripts (Figure 3A, Table S2). To test if these RBP binding sites are required for neurite localization or soma retention, we mutated 125 potential RBP binding sites in 5841 native 3’UTR sequences (Methods). We then obtained measurements of neurite/soma enrichment for the wild-type sequence, along with between one and 25 variants of the same sequence in which all instances of one RBP binding motif are mutated. Comparing each mutated sequence to its wild-type sequence yielded a readout of the effect of mutation of a given motif on the localization potential. As we perform this sequence alteration in up to 973 native sequences in parallel, we obtain an average effect of motif deletion that is independent of the specific context. Mutation of neurite-enriched motifs often had a significant negative effect on neurite localization, while mutation of soma-enriched motifs did not lead to increased neurite localization (Figure 3B, Figure S6A). Comparing the effect of motif mutations between CAD and Neuro-2a cells showed correlation between the cell lines for the mutation of neurite-enriched motifs (Pearson r=0.37, p=0.013, Figure S6B), providing further evidence that these motifs are required for RNA localization in both cell lines. We validated the effect of motif deletion by performing smFISH on a pair of otherwise identical sequences, one wild-type and one mutant (UCUUCU replaced by random sequences). In our MPRA, the wild-type sequence exhibits significant enrichment in the neurite fraction (log2FC=1.22 (CAD, p=0.00034) and 0.97 (Neuro-2a, p=0.0049), respectively), which is abrogated by UCUUCU mutations (log2FC=-0.57 (CAD, p=0.12) and 0.38 (Neuro-2a, p=0.10), respectively). Accordingly, smFISH signal in distal vs. proximal neurites was significantly lower in the mutant compared to wild-type (Figure 3C).

**Figure 3.**
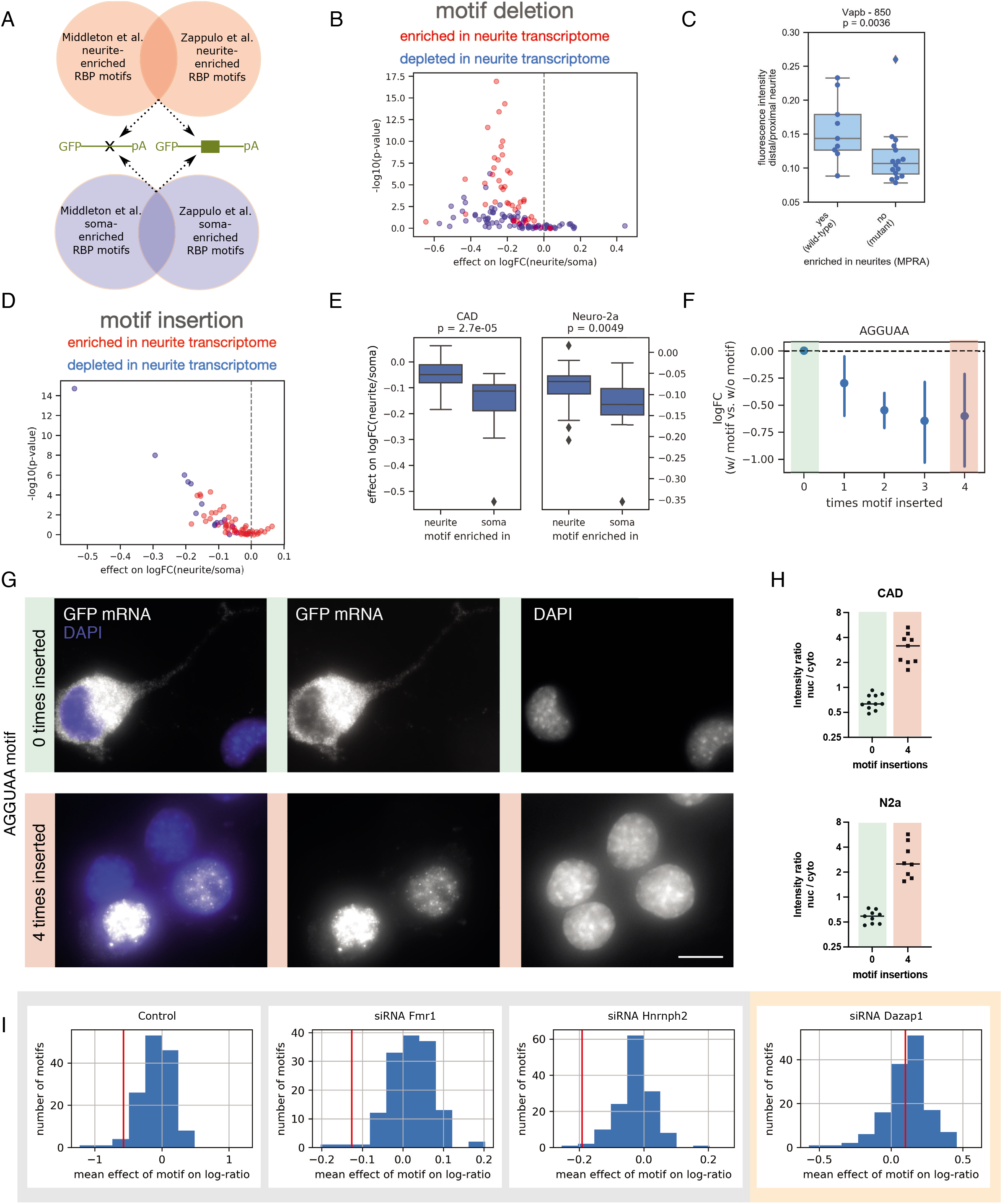
Single RBP motifs can only prevent, but not enforce neurite localization. A. Schematic of the bioinformatic analysis of available RNA-seq datasets to identify RBP motifs enriched in the soma or neurite transcriptome; these motifs were then either mutated in native 3’UTRs or inserted into native 3’UTR contexts. B. Each data point shows the mean effect on logFC(neurite/soma) in CAD cells for deletion of a specific motif in up to 973 native sequences, plotted against the associated p-value (Wilcoxon signed-rank test), for motifs identified as being enriched in neurite (red) or soma (blue) RNA-seq datasets (Middleton et al., Zappulo et al.). C. Quantification of dendritic localization (as determined by smFISH) for a pair of wild-type/mutant sequences: the wild-type sequence corresponds to positions 850-1000 in the Vapb 3’UTR, the mutant is the same sequence with all instances of the UCUUCU motif replaced by random sequences. D. Each data point shows the mean effect on logFC(neurite/soma) for insertion of a specific motif in up to 187 native sequences in CAD cells, plotted against the associated p-value (Wilcoxon signed-rank test), for motifs identified as being enriched in neurite (red) or soma (blue) RNA-seq datasets (Middleton et al., Zappulo et al.). E. Box plots showing the distribution of the mean effect of motif insertions on logFC(neurite/soma), for motifs identified as being enriched in neurite or soma RNA-seq datasets (Middleton et al., Zappulo et al.). F. Mean effect of the AGGUAA motif on logFC(neurite/soma) when inserted in a sequence 0-4 times. GH. smFISH images (G) and quantification of nucleus/cytoplasmic fluorescence intensity (H) for two individual 3’UTR reporters without (green) or with four AGGUAA motifs inserted (red, see corresponding MPRA data in panel F). I. Distribution of mean effect sizes of all motifs inserted in native 3’UTRs in control and different RNAi conditions; the red vertical line represents the mean effect of the AGGUAA motif in the different conditions.

To test if these RBP motifs can actively drive localization by themselves, we inserted 71 neurite- or soma-enriched RBP binding sites into up to 187 native contexts and, by comparing library sequences with or without the motif, measured their effect on the localization behavior of the transcript. In general, motif insertions could only shift localization towards soma, not towards neurite (Figure 3D, Figure S6C), with clear differences between motifs which we found in our analysis of endogenous RNA-seq data to be enriched in neurite and soma, respectively (Figure 3E; CAD: p=2.7×10^-5^, Neuro-2a: p=0.0049, Mann-Whitney-U test). Comparing motif effects between CAD and Neuro-2a cells (Figure S6B) showed a good correlation between the cell lines for the insertion of soma-enriched motifs (Pearson r=0.93, p=4.9×10^-7^), but not for neurite-enriched motifs (Pearson r=0.25, p=0.059), indicating that the effect of the soma-enriched motifs is robust between cell lines.

Some soma-enriched RBP motif insertions had highly significant effects on localization behavior, with AGGUAA showing the strongest negative effect, both in CAD (Figure 3D) as well as N2a cells (Figure S6D). This motif has been reported previously to be found preferentially in soma-enriched transcripts (Taliaferro et al., 2016). The effect increased with the number of binding sites introduced (Figure 3F). Comparing the distribution of neurite/soma enrichment of sequences with or without the AGGUAA motif introduced showed that this motif could not only prevent localization of the RNA to neurites, but often resulted in enrichment in the soma fraction (Figure S6E). To elucidate the mechanism by which AGGUAA leads to soma enrichment, we constructed individual reporter constructs of a sequence with or without four copies of the AGGUAA motif introduced. We performed smFISH as described above and observed a striking enrichment of the AGGUAA- containing reporter in the nucleus (Figure 3G,H).

AGGUAA is a potential binding site for the nucleocytoplasmic shuttling RBP Dazap1. To determine whether potential Dazap1 binding sites could more generally affect localization potential, we computed for every 3’UTR sequence tested in our MPRA a cumulative binding score for Dazap1 and >300 other RBPs based on position weight matrices obtained by RNAcompete (collected in ATtRACT, (Giudice et al., 2016)). We indeed observed a negative correlation between Dazap1 binding scores and logFC(neurite/soma) across all sequences tested (Pearson r=-0.085, p= 3.6×10^-77^; Spearman rho=-0.10, p=5.4×10^-107^). To further strengthen the link between Dazap1 and soma restriction, we downregulated Dazap1 (along with other RBPs whose binding sites were enriched in soma-restricted or neurite-localized genes, Figure 3A, Table S2). While the negative effect of introducing the Dazap1 motif on logFC(neurite/soma) was present in the control and other RNAi conditions, it was lost upon knock-down of Dazap1 (Figure 3I). This indicates that nuclear retention mediated by Dazap1 is one mechanism underlying soma enrichment and exclusion from neurites.

### Synthetic sequences can drive RNA localization to neurites

Our results on insertion or deletion of known RBP motifs suggest that a single motif cannot drive localization to neurites by itself. We therefore aimed to build and test longer synthetic 3’UTR sequences. We scanned 3’UTRs of neurite-localized and soma-restricted RNAs identified by Middleton et al. and Zappulo et al. for *de novo* motifs (Figure 4A). We used MEME Suite to discover novel, ungapped motifs in each of the sets that were enriched over the complementary compartment (i.e. neurite localized transcripts vs. soma localized transcripts). We then chose the best possible matches of the 20 top hits (for each compartment), omitting those with long homopolymeric stretches that would pose a problem for synthesis and subsequent steps. The remaining 66 synthetic sequences were then introduced in 69 native contexts and their distribution between neurite and soma fractions determined together with the rest of the library reporters. Unlike in the case of single RBP motifs whose introduction to a 3’UTR could only promote soma restriction, here we also identified longer synthetic sequences that resulted in increased enrichment in the neurite fraction (Figure 4B). The localization behavior of these synthetic sequences was similar in both cell lines used (Figure 4C, Pearson r=0.55, p=9.9×10^-7^). One of the synthetic sequences showed particularly robust localization to neurites, irrespective of the sequence context (Figure 4D, synthetic sequence 1). In other cases, the sequence context seemed to affect the localization behavior more; here, only a subset of insertion events led to efficient dendritic localization of the sequence containing the synthetic motif (Figure 4D, synthetic sequence 18).

**Figure 4.**
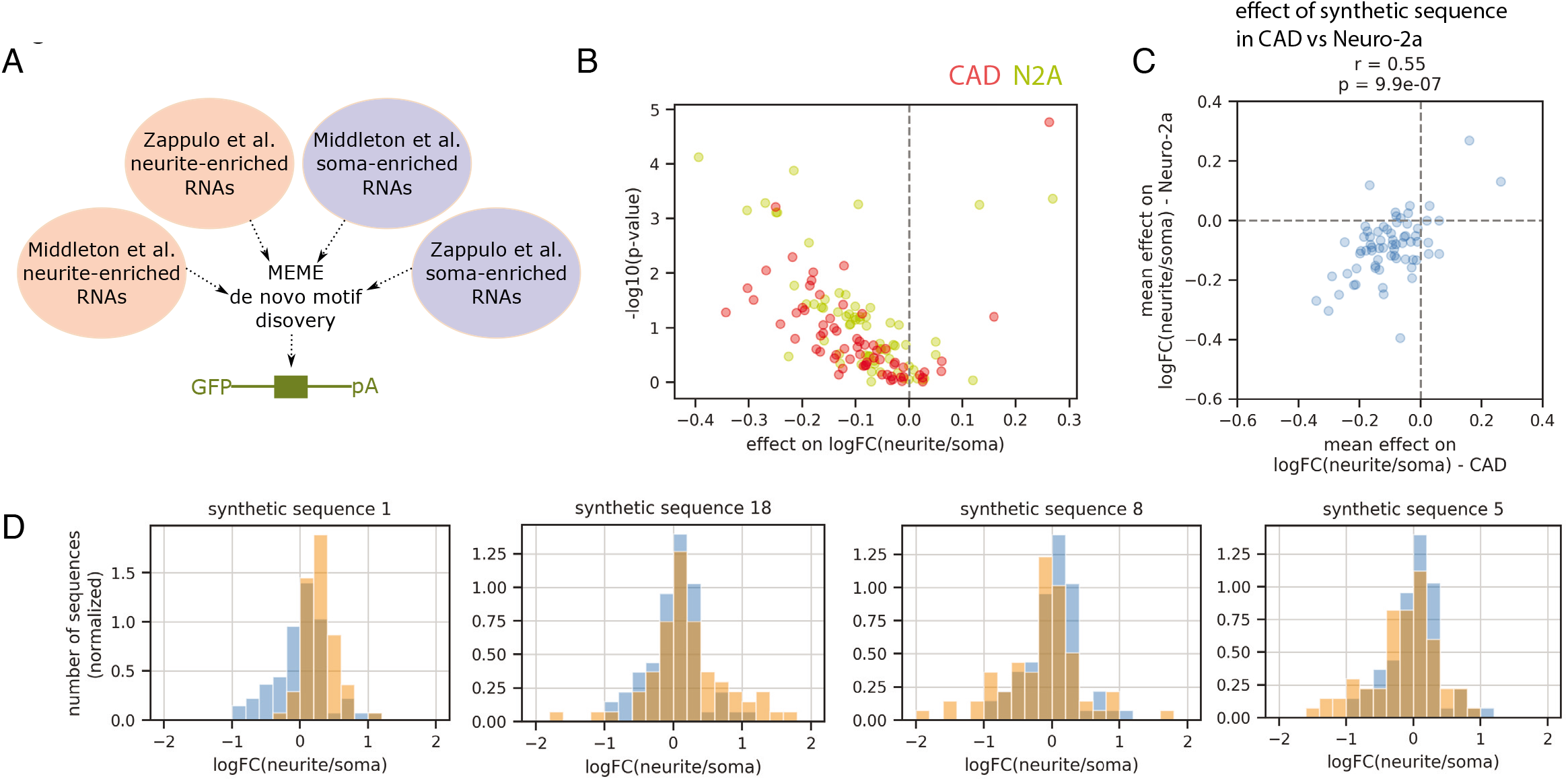
Synthetic sequences can drive RNA localization to neurites. A. Schematic of the bioinformatic analysis of available RNA-seq datasets to identify de novo motifs enriched in the soma or neurite transcriptome; these motifs were then inserted into native 3’UTR contexts. B. Each data point shows the mean effect on logFC(neurite/soma) for insertion of a synthetic de novo motif sequence in up to 69 native sequences, plotted against the associated p-value (Wilcoxon signed-rank test), in CAD (red) and Neuro-2a (yellow) cells. C. The mean effect on logFC(neurite/soma) for insertion of a synthetic de novo motif sequence in CAD cells is plotted against the effect of the same motif in Neuro-2a cells. D. Histogram of native 3’UTR sequences without (blue) or with (orange) insertion of the indicated synthetic sequence.

### Dissecting the mechanisms of neurite localization of a synthetic 3’UTR

To investigate by which mechanism synthetic 3’UTRs can localize to neurites, we selected a 97 nt long synthetic motif that exhibited strong neurite localization potential (synthetic UTR sequence 1, SU1; Figure 4D, left; Table S3) for biochemical analyses. As a control, we selected a pool of four sequences that were tested in our MPRA and did not show enrichment in neurites (Table S3). Both the SU1 and the control sequence pool were PCR-amplified with a T7 promoter sequence, *in vitro* transcribed, biotinylated, and used as baits for pulldowns against total protein lysates from CAD and Neuro-2a cells (Figure 5A). We analyzed the proteomic composition with liquid chromatography and tandem mass spectrometry (LC-MS/MS, methods) and detected a total number of 2172 proteins in CAD cell eluates and 2038 proteins in Neuro-2a cells, respectively. The resulting proteomic measurements of four SU1 eluates clustered separately from four control eluates in a heatmap of spearman correlations between all samples (Figure 5B). This indicates that the variation within replicate groups is considerably smaller than the observed variation of interest across groups. We performed differential protein expression analysis with Maxquant between the SU1 and the control groups, separately for each cell line (Figure 5C, Figure S7). In CAD cells, 549 proteins exhibited a differential expression between SU1 and control groups with an adjusted p-value <0.05 (Table S4). In Neuro-2a cells, 706 proteins were differentially detected between the groups (Table S5). We subjected the positively enriched proteins that were detected on the SU1 samples in both cellular contexts to protein list profiling with gProfiler 2 (Kolberg et al., 2020) and obtained several strongly enriched terms that reflect RNA binding and processing (Figure 5D).

**Figure 5.**
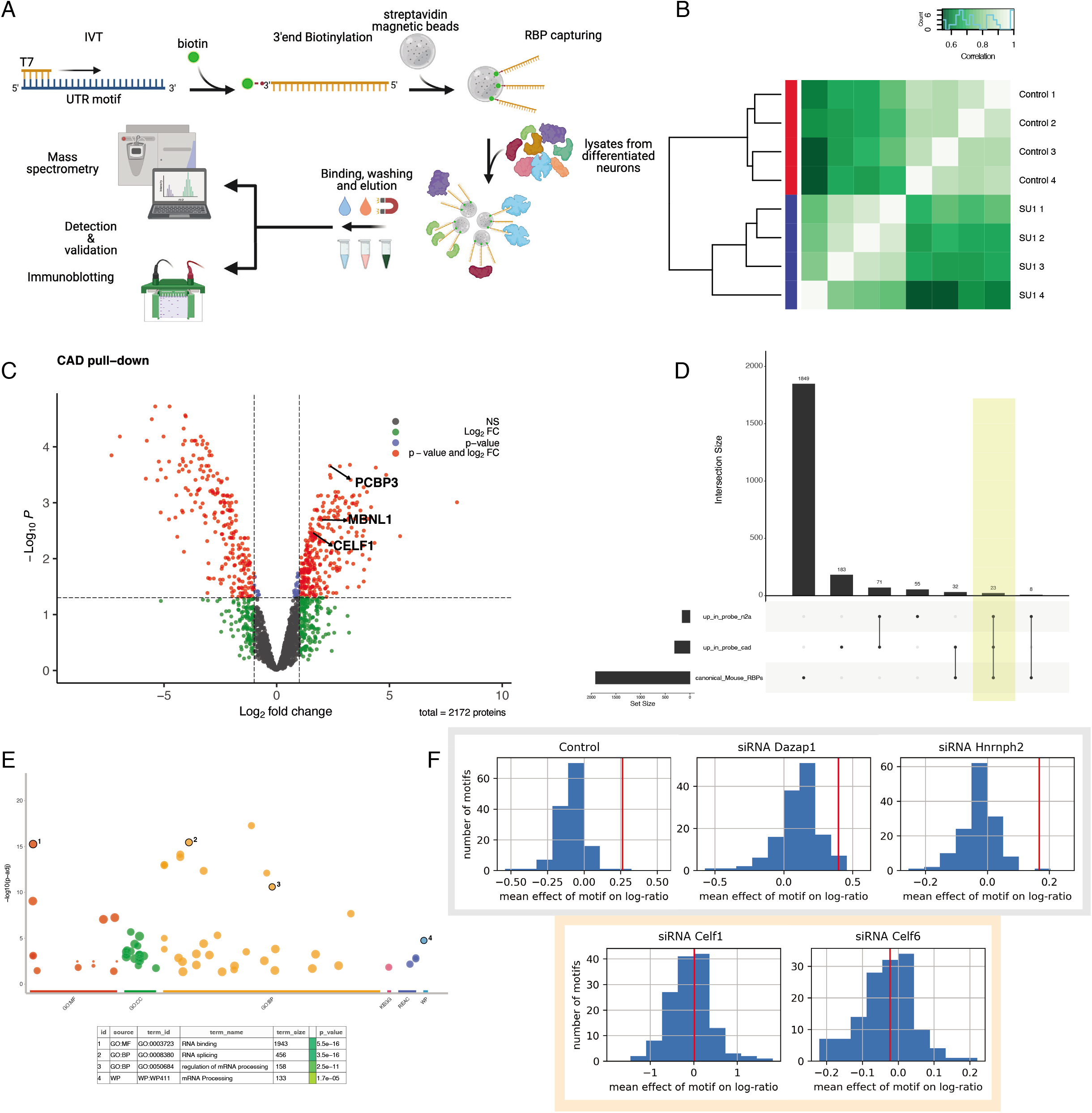
Mass spectrometric analysis reveals proteins that bind to the SU1 synthetic transcript and mediate its neurite localization potential. A. Schematic of the SU1 in vitro transcription and pulldown experiments for mass spectrometric analyses. B: Heatmap of across-sample spearman correlations in CAD cells. C: Differential protein pulldown analysis in CAD cells. D: Gprofiler 2 enrichment analyses of the intersection of significantly pulled-down proteins with the SU1 probe in CAD and N2A cells; gene sets of particular interest are highlighted in the legend. E: Quantitative intersection analysis of positively and negatively enriched proteins across both cell lines and an annotated list of canonical mouse RBPs (Liao et al., 2020). F. Distribution of mean effect sizes of all motifs inserted in native 3’UTRs in control and different RNAi conditions; the red vertical line represents the mean effect of the SU1 motif in the different conditions.

We then set out to identify RNA binding proteins that could mediate the observed neurite localization pattern of SU1 in our proteomic dataset. To this end, we intersected the significant positively enriched proteins from both cell lines with a database of annotated RBPs (Liao et al., 2020) and retrieved a set of 24 RBP candidates for further investigation (Figure 5E). We complemented the biochemical study of SU1 binders with an *in silico* analysis of RBP motif presence using RBPmap (Paz et al., 2014) (Table S6). Several RBP candidates from our intersection analysis (Figure 5E) exhibited a significant binding motif enrichment in the SU1 sequence: Celf1 (z-score 2.643, p=4.11×10^-3^), Mbnl1 (z-score 1.974, p=2.42×10^-2^), and Rbm38 (z-score 4.333, p=7.35×10^-6^). The combined evidence from biochemical pulldowns and binding motif analyses led us to choose Celf1 for functional validation with RNA interference assays. Additionally, we abrogated Celf6, an RBP that was predicted to bind to the same SU1 sequence stretch as Celf1 by RBPmap. The abrogation of both Celf1 and Celf6 led to a loss of the localization potential of SU1 when compared to non-targeting control, Dazap1, or Hnrnph2 RNAi experiments (Figure 5F), indicating that these proteins are necessary for SU1 subcellular localization.

### MPRA-trained models predict subcellular localization of endogenous 3’UTRs

Our large dataset of neurite/soma distributions of 3’UTR reporter sequences provides a starting point for deciphering the regulatory logic of RNA localization and for prediction of the localization behavior of novel sequences. Since our data suggested that both neurite as well as soma enrichment can be actively mediated by sequence motifs, we trained two classifiers on 90% of our data (XGBoost, gradient boosting decision trees, see Methods): One to discriminate between sequences driving significant neurite enrichment (p<0.05) and all others and one to discriminate between sequences driving significant soma enrichment (p<0.05) and all others (Figure 6A). The remaining 10% served as the test set and were not used at any point in building the model.

**Figure 6.**
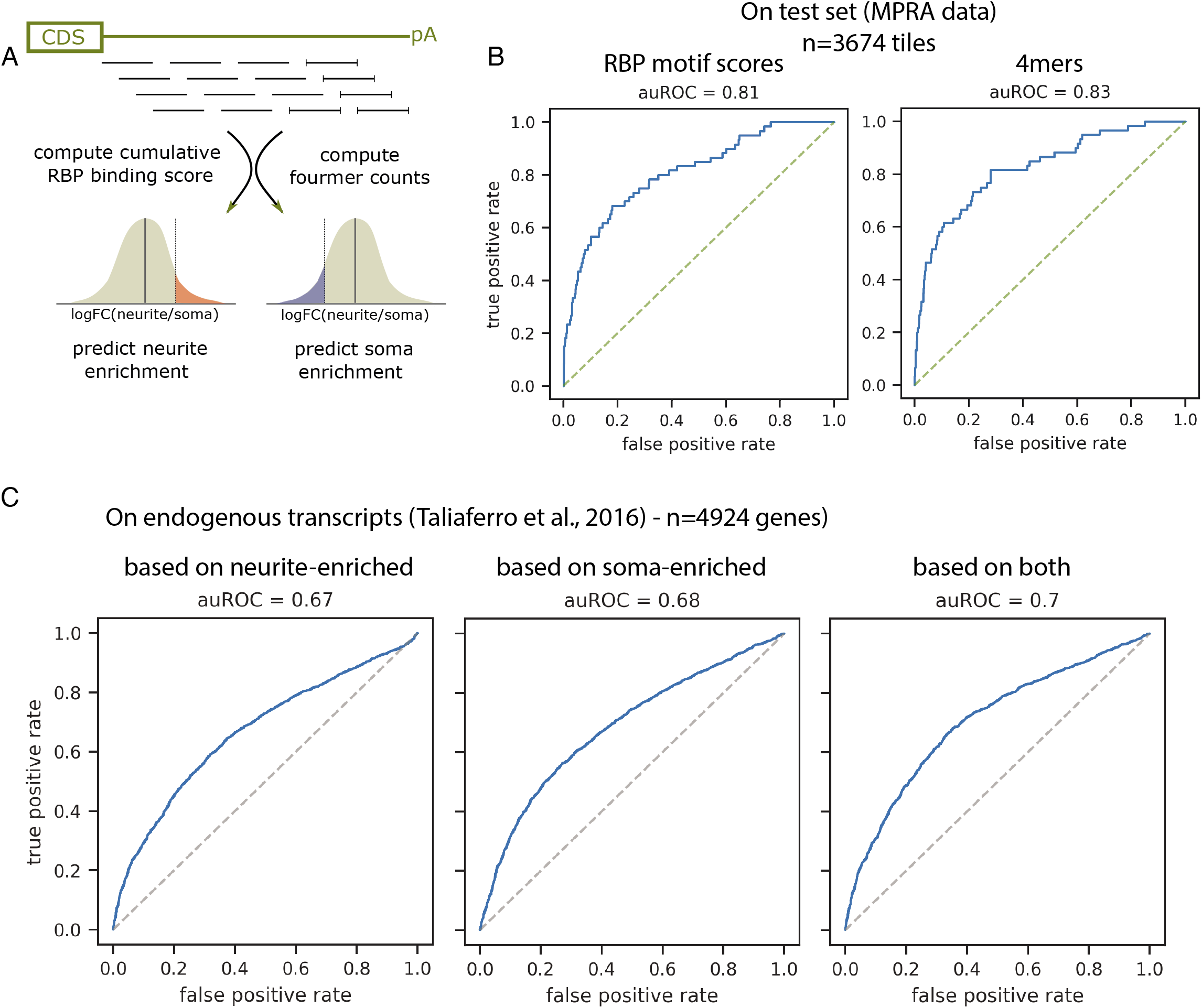
MPRA-trained models predict subcellular localization of endogenous 3’UTRs. A. Outline of the prediction strategy. B. Receiver operating characteristic (ROC) curves showing performance of a classifier trained on 90% of all library sequences passing filtering and testing on the remaining 10% (n=3674; held-out test data), using cumulative RBP motif scores or 4mer counts as features; auROC: area under the ROC curve. C. Receiver operating characteristic (ROC) curves showing performance of a classifier trained on 90% of all library sequences passing filtering and testing on 4924 genes for which neurite/soma distribution in CAD cells has been determined previously (Taliaferro et al., 2016); the prediction constitutes the combined prediction for 150mers from the native 3’UTR, based on models predicting neurite enrichment (left), soma enrichment (middle) or the combined output of both (right).

RBPs are thought to be the trans acting factors mediating localization to neurites. We therefore computed cumulative binding scores for a set of 218 RBPs for which binding sites have been identified (RNAcompete, (Ray et al., 2017)) and used these as features to train our models. As an alternative, unbiased approach, we used counts of all possible 4mers in the sequence as features. We scored performance of prediction algorithms and parameter settings on the training set by cross-validation and then trained both predictors on the entire training set (33,057 sequences). The combined output of both models was able to predict the localization behavior of unseen variants with high accuracy (Figure 6B; area under the receiver operating characteristics curve (auROC) was 0.81 for motif scores and 0.83 for 4mers). 4mers showed a slightly better performance, indicating that restricting the features to known RBP binding sites might not capture all the relevant information. We used Shapely (SHAP) values (Lundberg et al., 2020) for determining the contribution of each feature to the prediction result of every sample (Figure S8). This identified the sequence elements and potential RBP binding sites driving the prediction, highlighting links to potential trans-acting factors mediating the neurite or soma enrichment.

According to our data, the localization potential of a native 3’UTR tends to be broadly encoded in the sequence and is not necessarily restricted to one clearly defined localization motif. In order to apply our model to predict the localization behavior of native 3’UTRs we therefore chose to first predict the localization potential of individual tiles of native 3’UTRs, defined the same way as in our library (150 nt in length, 50 nt step size between tiles). For each of these tiles our models predict the likelihood that it will have a significant effect on neurite and soma localization, respectively. In line with our observation that the existence of a soma-enriched segment in a 3’UTR sequence can have a dominant-negative effect and prevent localization of the transcript, prediction of neurite localization based on (lack of) somatic enrichment of any tile performed as well as prediction based on a significant dendritic enrichment as the positive class (auROC=0.68 vs auROC=0.67; Figure 6C). Combining both prediction strategies slightly improved prediction further (auROC=0.7). These results indicate that the localization potential of a native 3’UTR can be inferred by estimating contributions of different segments individually.

## Discussion

Mechanism and functional importance of RNA localization have been central topics in biological research over the last decades. Despite numerous insights into the localization of well characterized transcripts, a general understanding of the link between sequence and function is still missing, curbing our ability to predict the effect of sequence alterations on localization dynamics.

Here, we established a novel experimental approach that enables the mapping of RNA sequence motifs to subcellular localization. The systematic nature of our assay and the large number of sequences tested allowed us to investigate the general principles and bring us one step closer to deciphering the sequence-encoded rules of RNA localization.

Our results agree with the existing literature about known 3’UTR-regions that encode neurite- localization potential; Tushev et al. have shown that the dendritic localization potential of Camk2a is encoded mostly in its longest isoform (Tushev et al., 2018), and our strongest neurite- localization potential lies within the gene region that is specific to the long isoform.

From our measurements of the localization behavior of 13,753 native sequences and 34,236 designed sequence variants the following model emerges: We propose that in most cases the localization potential is broadly encoded along the length of the 3’UTR sequence. While in some genes a defined region with strong potential for neurite localization can be identified, this is not the case for many other transcripts found in neurites. While we cannot rule out that for some of these genes our MPRA is not an adequate tool to identify the localization regions, our results show that the collective localization behavior of all the 3’UTR tiles combined does recapitulate the localization reported for the endogenous transcript. We therefore suggest that many small contributions, e.g. RBP binding events, can slightly bias localization of a transcript towards neurite or soma (Figure 7A,B). All these small contributions combined can then drive neurite localization of the entire transcript. This way of encoding neurite localization might be more robust to small sequence changes and more effective in preventing ectopic localization of soma-restricted transcripts. This model is in line with the fact that despite large experimental efforts over the last decades only a small number of focused localization elements could be identified.

**Figure 7.**
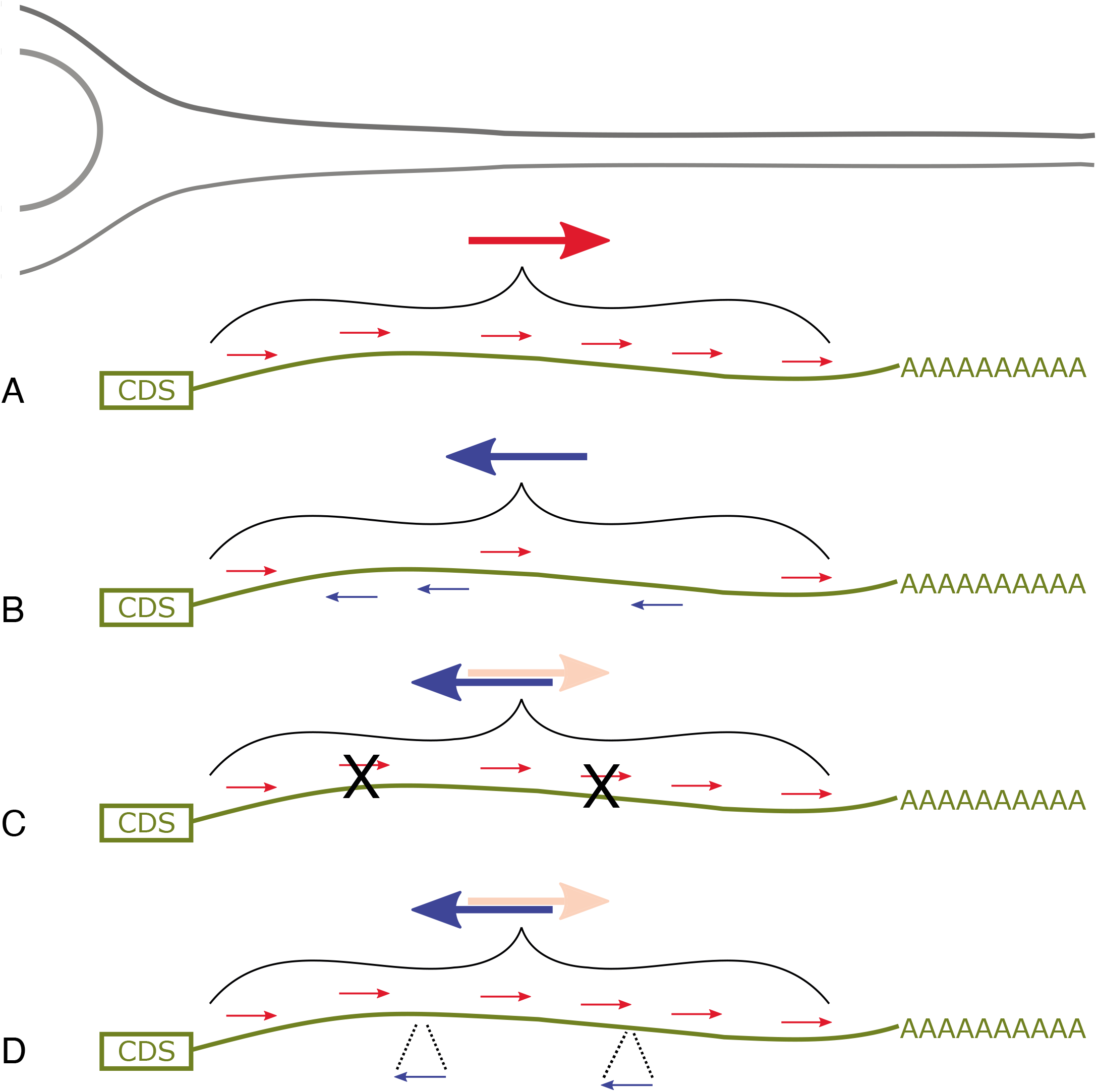
Localization potential as a sum of small contributions. A. Sequence elements promoting neurite localization (small red arrows) encoded along the length of the 3’UTR promote enrichment of a transcript to neurites (large red arrow). B. Contributions from sequence elements promoting neurite (small red arrows) and soma (small blue arrows) neutralize each other, leading to restriction of the transcript to the soma (large blue arrow). C. Mutation (“X”) of sequence elements promoting neurite localization (small red arrows) can abolish enrichment of the transcript in neurites. D. Introduction of a sequence element promoting soma restriction (small blue arrow) can prevent localization to neurites.

Based on these principles, we developed a computational model to predict the localization behavior of a native 3’UTR based on the contribution of small parts. The model trained on our MPRA data was indeed able to discriminate between neurite-localizing and non-localizing endogenous transcripts with good accuracy, corroborating our view of the localization behavior as a sum of different contributions from 3’UTR tiles promoting neurite or soma localization.

At the level of individual 3’UTR tiles, localization to neurites can be abrogated by mutating individual RBP binding motifs (Figure 7C), but it cannot be created by introducing potential RBP binding sites. The preferences for functional binding of an RBP are probably more complex and go beyond the narrow sequence motif. Therefore merely introducing an RBP binding motif is not sufficient to actively drive translocation of a transcript to neurites.

In contrast, enrichment of an mRNA in neurites can be actively prevented by introducing sequence motifs (Figure 7D). Our results reveal one possible mechanism by which this observation can be explained mechanistically, namely nuclear retention. We identify a potential Dazap1 binding motif as a promoter of nuclear retention and consequently as a strong inhibitor of neurite enrichment. This Dazap1-mediated nuclear retention presented the most drastic effect in our assay, and we postulate that other associations with soma-restricted RBPs underlie our observation of widespread dominant negative activity of soma-enriched motifs on neurite localization.

Since the context-dependent addition of short RBP motifs did not suffice in encoding neurite- localization potential, we subsequently assembled larger de-novo motifs based on observed consensus sequences across endogenous 3’UTRs. We took a synthetic biology approach and attempted to understand localization mechanisms by building a localizing synthetic 3’UTR *de novo*. This approach indeed yielded sequences which robustly localized to neurites. We then dissected the localization potential of one of our synthetic 3’UTR sequences. Converging evidence from bioinformatic analysis, mass spectrometry and RNAi experiments pointed at a number of prominent RBPs which have not been directly implicated in RNA localization up to date, in particular Celf1 and Celf6. These results show that high-throughput functional testing can reveal novel players of RNA localization and advance our understanding of the complex interplay between RBPs and the transcriptome.

Our experimental strategy allows us to perform high-throughput testing of the localization potential of a sequence, but entails also a number of trade-offs: As efficient transfection of the large number of reporter constructs is key to obtaining high-quality quantitative data, we chose two neuronal cell lines, CAD and Neuro-2a cells, as model systems. In vivo, localization of certain RNAs can depend on cell identity, neural activity or the tissue context. While these specific features of RNA localization cannot be fully reproduced in a neuronal cell line, we believe that the general principles of RNA localization to neurites will be similar, and predictions from our MPRA can subsequently be tested in a targeted way in primary neurons or other suitable models that are incompatible with the large scale of our MPRA library. In addition, the synthesis of a large number of rationally designed sequences limits the length of the sequence that can be tested (to 150 nt in our case). While some more complex localization motifs might be missed by our current MPRA setup, our data show that we can detect many known and novel localization motifs across transcripts. Furthermore we show that taken together, the 3’UTR tiles tested in our assay recapitulate the endogenous localization behavior of the full transcript.

Many neurological diseases have been linked to dysregulation of RNA localization (Wang et al., 2016). In the future, the high-throughput approach developed here can enable us to predict and experimentally test how genetic variation affects the localization potential of a sequence. This can reveal the functional consequence of disease-associated genetic variants and thereby highlight new therapeutic strategies. Beyond the model system employed here, this approach can be extended to the study of subcellular RNA localization in other tissues like intestinal epithelia, which will allow for a systematic comparison of regulatory mechanisms of RNA localization.

## Supplementary Figure Legends

Figure S1.

Bright field images of CAD cells grown in differentiation medium (day 6).

Figure S2.

Measurements (upper panel: logFC(neurite/soma), lower panel: -log_10_(p-value) of the effect, multiplied by the sign of the logFC) for tiles along the 3’UTR of genes with a region of increased neurite localization potential, as measured in CAD (red) and Neuro-2a (yellow) cells; gray denotes the area with p>0.05.

Figure S3.

Measurements (upper panel: logFC(neurite/soma), lower panel: -log_10_(p-value) of the effect, multiplied by the sign of the logFC) for tiles along the 3’UTR of genes with broadly encoded localization potential, as measured in CAD (red) and Neuro-2a (yellow) cells; gray denotes the area with p>0.05.

Figure S4.

A. Each data point represents the mean logFC(neurite/soma) of all tiles corresponding to a segment taken from the same native 3’UTR (left) or the mean logFC of the three tiles with the highest statistical significance of enrichment in any compartment (right); the groups correspond the overlap of the 30 genes with the highest neurite (“top neurite enriched RNAs”) or soma enrichment (“top soma enriched RNAs”) in both CAD and Neuro-2a cells as measured by Taliaferro et al. B. logFC(neurite/soma) measured in CAD (red) and Neuro-2a (yellow) cells are plotted against the p-value of the enrichment, for genes with a defined region of increased neurite localization potential (left), genes with broadly encoded localization potential (middle) or genes with no evidence for neurite localization in endogenous RNA-seq data (right).

Figure S5.

Measurements (upper panel: logFC(neurite/soma), lower panel: -log_10_(p-value) of the effect, multiplied by the sign of the logFC) for tiles along the 3’UTR of genes with no evidence for neurite localization in endogenous RNA-seq data, as measured in CAD (red) and Neuro-2a (yellow) cells; gray denotes the area with p>0.05.

Figure S6.

A. Each data point shows the mean effect on logFC(neurite/soma) for deletion of a specific motif in Neuro-2a cells in up to 973 native sequences, plotted against the associated p-value (Wilcoxon signed-rank test), for motifs identified as being enriched in neurite (red) or soma (blue) RNA-seq datasets (Middleton et al., Zappulo et al.). B. The mean effect on logFC(neurite/soma) for mutation (top) or insertion (bottom) of an RBP motif (neurite-enriched, soma-enriched or both) in CAD cells is plotted against the effect of the same motif in Neuro-2a cells. C. Each data point shows the mean effect on logFC(neurite/soma) for insertion of a specific motif in up to 187 native sequences in Neuro-2a cells, plotted against the associated p-value (Wilcoxon signed-rank test), for motifs identified as being enriched in neurite (red) or soma (blue) RNA-seq datasets (Middleton et al., Zappulo et al.). D. Histogram of native 3’UTR sequences without (blue) or with (orange) insertion of the AGGUAA motif in CAD (left) and Neuro-2a cells (right).

Figure S7.

A: Heatmap of across-sample spearman correlations in N2A cells. B: Differential protein pulldown analysis in N2A cells.

Figure S8.

Features driving prediction of neurite or soma enrichment. Effects (as determined using SHAP) of 4mers (top) and cumulative RBP binding scores (bottom) on the model prediction (ranked by their importance for the prediction) for a classifier built on the indicated feature sets and predicting significant neurite (left) or soma (right) enrichment (p<0.05). The color denotes the feature value and the position along the x-axis denotes the impact on model output, for each item in the training set.

## Supplementary Tables

Table S1. List of genes used in the library design.

Table S2. Motif enrichment analysis on published neurite and soma transcriptome datasets.

Table S3. Sequences of a synthetic neurite-enriched sequence (SU1) and four soma-restricted library sequences used in the pull-down experiments.

Table S4. Mass spectrometry results (SU1 vs. soma restricted sequences), CAD cells.

Table S5. Mass spectrometry results (SU1 vs. soma restricted sequences), Neuro-2a cells.

Table S6. RBP motifs present in the SU1 sequence (RBPmap).

Table S7. Sequence of the smFISH probes used to detect the *gfp* coding sequence.

## Methods

### Synthetic library design

Oligonucleotides were designed to maintain a constant length of 198 nt. Restriction sites used for cloning were excluded from the design. All the variants were composed of an 18 nt forward primer, 12 nt barcode sequence, 150 nt variable region and 18 nt reverse primer sequences. DNA barcodes were designed to differ from any other barcode in the library in at least 3 nt.

Design of the subsets of the library was carried out in Python.

#### Tiling of endogenous 3’UTRs

Tiles of 150 nt length were chosen, starting from the first position after the stop codon and extending until the most distal poly-adenylation site, excluding tiles that would contain a poly-adenylation site themselves to avoid cleavage within the 3’UTR reporter construct. Genes were selected based on previous transcriptomics data obtained from cultured neurons (Ciolli Mattioli et al., 2019; Middleton et al., 2019; Taliaferro et al., 2016; Zappulo et al., 2017). These consisted of the following groups of genes (see also Table S1): 14 RNAs identified in multiple prior studies (e.g. Camk2a, Map2, Shank1), yielding 715 tiles; the 40 RNAs found in most often in single dendrites analyzed by (Middleton et al., 2019), yielding 1178 tiles; 347 dendrite enriched genes found by Middleton et al., yielding 7948 tiles; 127 neurite enriched (logFC>2, adj. p-value<0.01) genes identified by (Zappulo et al., 2017), yielding 2355 tiles; 33 genes with evidence from two studies (Ciolli Mattioli et al., 2019; Middleton et al., 2019) for differential localization behavior of 3’UTR isoforms, yielding 1203 tiles; 5 genes found enriched in the soma fraction in all of the previous studies (Taliaferro, Zappulo, Middleton, Ciolli Matolli et al., 2018), yielding 710 tiles. After removing duplicate sequences and sequences containing potential poly-adenylation sites, the final library covering endogenous 3’UTRs consisted of 13754 tiles.

#### Multiple barcode controls

We added multiple variants to the library that contained the same variable region, but different barcodes, in order to gauge potential effects of the barcode and the technical noise of our assay.

#### RBP motif insertions and deletions

We assembled a list of potential RBP binding sites consisting of the results of our own bioinformatic analysis (see details below: RBP motif enrichment analyses) and previously reported motifs (neurite-enriched according to Middleton et al.: ATCAACG, ATCATCG, TTCGAT, CCGCAA, GTGGGT; neurite-enriched according to Taliaferro et al.: GCTGCT, CTGCTG, GCGCTG, CTGGAC, CCTGCT, TCTGGA, CCCCAA, CTGCCC, ACACTG, TTTTCA, TTTTTT, ATACAG; soma-enriched according to Taliaferro et al.: TAGGTC, TCTTCT, CTCTTT, TCTCTT, TCTCTC, AGGTAA).

#### Motif mutations

We mutated 44 neurite- and 84 soma-enriched motifs in endogenous 3’UTR tiles (see above) in the following way: For genes with evidence for neurite enrichments (all groups of genes mentioned above with the exception of the five genes found enriched in the soma compartment in all of the previous studies, we scanned each tile for the presence of any of the 44 neurite-enriched motifs. If a motif was present, we included a sequence in the library which corresponded to the endogenous 3’UTR tile, but had all instances of this motif mutated (replaced by a random sequence). For genes previously found enriched in the soma compartment and for those genes with differential localization behavior of 3’UTR isoforms we scanned each tile for the presence of any of the 84 soma-enriched motifs. If a motif was present, we included a sequence in the library which corresponded to the endogenous 3’UTR tile, but had all instances of this motif mutated (replaced by a random sequence).

#### Motif insertions

We introduced these motifs in different configurations:

We inserted two copies of 21 neurite- and 15 soma-enriched motifs in 189 native contexts (150 nt tiles from endogenous 3’UTRs) at positions 50 and 100 in order to test the activity of the motifs in as many native contexts as possible.

We inserted one, two, three or four copies of 21 neurite- and 15 soma-enriched motifs in 22 native contexts (150 nt tiles from endogenous 3’UTRs) at positions 30, 60, 90 and 120 in order to determine if the effect of the motif increases with the number of times it is present.

We inserted one or two copies of 44 neurite-enriched motifs in 22 native contexts (150 nt tiles from endogenous 3’UTRs) at position 50 or 50 and 100, in the native sequence or embedded within an artificial 9 bp hairpin structure to determine the effect of the local secondary structure on motif effects.

We inserted 66 de novo synthetic motifs (see below) in 69 native contexts (150 nt tiles from endogenous 3’UTRs) at position 30.

### Synthetic library cloning

The cloning steps were performed essentially as described previously (Vainberg Slutskin et al., 2018; Mikl et al., 2019). We obtained the oligonucleotide library from Twist Bioscience as a pool. The two subsets of this pool corresponding to native 3’UTR tiles and designed sequence alterations (mutations and motif insertions) were defined by unique amplification primers. The oligo pool was resuspended in 10 mM Tris buffer, pH 8.0 to a concentration of 20 ng/μl. We amplified both libraries by performing 4 PCR reactions, each of which contained 19 μl of water, 1 μl of the oligo pool, 10 μl of 5× Herculase II reaction buffer, 5 μl of 2.5 mM deoxynucleotide triphosphate (dNTPs) each, 5 μl of 10 μM forward primer, 5 μl of 10 μM reverse primer, and 1 μl Herculase II fusion DNA polymerase (Agilent Technologies). The parameters for PCR were 95°C for 1 min, 14 cycles of 95°C for 20 s, and 68°C for 1 min, each, and finally one cycle of 68°C for 4 min. The oligonucleotides were amplified using library-specific common primers, which have 18-nt complementary sequence to the single-stranded 198-mers and a tail containing SgsI (forward primer) and Sdal (reverse primer) restriction sites (native 3’UTR tiles library: cacaGGCGCGCCaCGAAATGGGCCGCATTGC and cacaCCTGCAGGaTCGTCATCAGCCGCAGTG; designed sequence alterations library: cacaGGCGCGCCaGACAGATGCGCCGTGGAT and cacaCCTGCAGGaGCATTGGATCGGGTGGCT. The PCR products were concentrated using Amicon Ultra, 0.5 ml 30K centrifugal filters (Merck Millipore). The concentrated DNA was then purified using PCR mini-elute purification kit (Qiagen) according to the manufacturer’s protocol. Purified library DNA was cut with the unique restriction enzymes SgsI and SdaI (Fermentas FastDigest) for 2 hours at 37°C in two 40-μl reactions containing 4 μl fast digest (FD) buffer, 1 μl SgsI enzyme, 1 μl Sdal enzyme, 18 μl DNA and 16 μl water, followed by heat inactivation for 20 min at 65°C. Digested DNA was purified, first using PCR mini-elute purification kit (Qiagen) and then using 2.2x SPRI beads (Beckman-Coulter).

The master plasmid for inserting the library was created by introducing a synthetic sequence containing a stop codon, a primer binding site and restriction sites for SgsI and SdaI into the Bsp1704I site at the 3’ end of the GFP coding region of pcDNA3-EGFP (Addgene #13013). The modified plasmid was cut with SgsI and SdaI (Fermentas FastDigest) in a reaction mixture containing 6 μl FD buffer, 3 μl of each enzyme and 3.5 μg of the plasmid in a total volume of 60 μl. After incubation for 2.5 hours at 37°C, 3 μl FD buffer, 3 μl alkaline phosphatase (Fermentas) and 24 μl water were added and the reactions were incubated for an additional 30 mins at 37°C followed by 20 min at 65°C. Digested DNA was purified using a PCR purification kit (Qiagen). The digested plasmid and DNA library were ligated for 30 min at room temperature in 15 μl reactions, containing 150 ng plasmid and the insert in a molar ratio of 1:1, 1.5 μl FastLink 10× ligation buffer, 1.5 μl ATP and 1 μl FastLink DNA ligase (Lucigen Corporation), followed by heat inactivation for 15 min at 70°C. Ligated DNA was transformed into *E. cloni* 10G SUPREME Electrocompetent Cells (Lucigen) (2 μl of the ligation mix per reaction) using Biorad GenePulser Xcell (Voltage 1800V, capacitance 25 uF, resistance 200 Ohm, 1 mm cuvettes), which were then plated on 4 Luria broth (LB) agar (200 mg/ml amp) 15-cm plates per transformation reaction (25 μl). The rationally designed (12,809 variants) and the native (5000 variants) parts of the library were cloned separately. For the two libraries we collected around 1.2×10^6^ and 2.3×10^6^ colonies, respectively, the day after transformation by scraping the plates into LB medium. Library-pooled plasmids were purified using a NucleoBond Xtra EndoFree midi prep kit (Macherey Nagel). To ensure that the collected plasmids contain only a single insert of the right size, we performed colony PCR (at least 30 random colonies per library).

### Cell culture

CAD cells were acquired from Sigma-Aldrich (#ATCC^®^ CRL-11179); Neuro-2a cells were a gift from Prof. Peter Scheiffele, U. Basel, Switzerland. CAD cells were grown in DMEM/F-12 (Gibco) supplemented with 8% fetal bovine serum (Gibco, #10270-106) and 1% Penicillin-Streptomycin solution (Gibco, #15140-22). Neuro-2a cells were grown in DMEM (Gibco) supplemented with 10% fetal bovine serum and 1% Penicillin-Streptomycin solution. The cells were grown in a humified incubator at 37°C and 5% CO2 and split by dissociation by pipetting (CAD cells) or by trypsinizing (TrypLE, ThermoFisher). To induce differentiation into a more neuron-like phenotype, medium was changed to differentiation medium (DMEM/F-12 with 0.8% fetal bovine serum and 1% Penicillin-Streptomycin solution and DMEM with 1% fetal bovine serum and 1% Penicillin-Streptomycin solution, respectively).

### Library transfection and neurite- and soma-specific RNA extraction

For quantifying the abundance of library sequences in neurite and soma, CAD and Neuro-2a cells were grown on Millicell Hanging Cell Culture Inserts with a pore size of 3 μm (Millipore #MCSP06H48), adapting experimental pipelines described earlier (Ludwik et al., 2019; Taliaferro et al., 2016; Zappulo et al., 2017). Prior to seeding the cells, the bottom of the insert was coated with 100 ul Matrigel (Corning #356237, diluted to 3 mg/ml with PBS). Matrigel was allowed to solidify for 3 hours by incubating the plates with the coated inserts bottom up at 37°C and 5% CO2. 5×10^5^ cells were seeded on each filter in differentiation medium, adding medium also to the bottom compartment of the well. After 24 hours, reporter libraries were transfected into the cells using 2.5 ug reporter DNA per well and Lipofectamine 3000 (ThermoFisher) according to the manufacturer’s protocol. For library transfections, three biological replicates were performed. Each replicate was pooled from three 6-well plate inserts with transfected cells. 24 hours after transfection soma and neurites were harvested as follows: Wash cells twice with PBS. Aspirate and add as much fresh PBS to wells with inserts as needed so that the volume above the filter insert will remain approximately 1 ml. Scrape off cells growing on top of the insert, wash off cell bodies with P1000 pipette and transfer them to a microcentrifuge tube. Spin down at 300 g for 5 min, take off supernatant and add 300 ul TRIreagent. While spinning down the cell bodies, wash the remaining inserts with PBS. Clean the insert filter thoroughly, scrape the insert to remove all the remaining cell bodies carefully without disrupting the filter, aspirate pbs. Repeat the step at least one more time and check under the microscope if all the cell bodies have been removed; this step is critical for good separation. Aspirate PBS, take out insert, remove filter from the insert with forceps and transfer it to a microcentrifuge tube with 300 ul TRIreagent.

RNA was isolated using Direct-zol RNA miniprep kit (Zymo research #R2051) according to the manufacturer’s protocol.

### cDNA synthesis and library preparation for Illumina sequencing

cDNA was synthesized from up to 10 ug total RNA in a 40 ul reaction using SuperScript IV Reverse Transcriptase (ThermoFisher) according to the manufacturer’s protocol, with oligo-dT primers containing 6 nt barcodes, a 15 nt unique molecular identifier (UMI) and a partial Illumina read 1 primer sequence. For library preparation, reporter cDNA was PCR amplified using a reporter specific forward primer and a reverse primer binding the anchor sequence of the oligo-dT primer (corresponding to the Illumina TruSeq Read 1 sequence):

20 ul Kapa hifi ready mix, 1.5 ul 10 uM primer reporter-specific forward primer (adding the Illumina TruSeq Read 2 sequence) and 1.5 ul reverse primer (Illumina TruSeq Read 1), 4 ul cDNA, 13 ul water (2 reactions per sample). 98 deg 3 min, 15 (soma) or 20 (neurite) cycles: 98 deg 20 sec, 65 deg 15 sec, 72 deg 20 sec; 72 deg 1 min. After cleaning up the reaction with 1.8x SPRI beads (Beckman-Coulter) Illumina sequencing adaptors (NEBNext Multiplex Oligos for Illumina, NEB #E7600S) were added in a second PCR reaction: 20 ul Kapa hifi ready mix, 1 ul i7 and 1 ul i5 (from NEB #E7600S), 8 ul 1st PCR product after SPRI beads, 10 ul water. PCR program: 98 deg 3 min, 12 cycles: 98 deg 20 sec, 65 deg 15 sec, 72 deg 20 sec; 72 deg 1 min. The reactions were cleaned up with SPRI beads (0.6 x) and the size of the product was verified using Tapestation (Agilent High Sensitivity D1000 ScreenTape).

### RNAi experiments

siRNA pools targeting mouse Dazap1, Celf1, Celf6 or Hnrnph2 were obtained as a siGENOME SMARTPool from Dharmacon. siRNA transfections were carried out 2 days before library transfections using Dharmafect 1 according to the manufacturer’s protocol. Target knockdown was verified using qRT-PCR.

### In vitro transcription and pull-down experiments

10uL (~20-35ng/uL) of cleaned-up PCR amplicon (primer sequences are provided in the table below, used with screening pooled library) were used as template of the in vitro transcription (HiScribe^™^ T7 Quick High Yield RNA Synthesis Kit; #E2050S, New England Biolabs), performed at 37°C for 16h, followed by DNAseI treatment (37°C for 15’). IVT RNAs were then cleaned-up and concentrated (DNA Clean & Concentrator-5; #D4013, Zymo Research)

3’-desthiobiotin labeling was carried following the manufacturers’ guidelines of Pierce^™^ RNA 3’ End Desthiobiotinylation (ThermoFisher, #20163). Briefly, ~115pmol of each RNA were first subjected to fast denaturation in the presence of 25% v/v DMSO (85°C for 4’) to relax 2nd structures, and subsequently labelled at 16°C for 16h. RNA binding proteins were isolated by the means of Pierce^™^ Magnetic RNA-Protein Pull-Down Kit (ThermoFisher, #20164). Briefly, 3’- desthiobiotin labelled RNAs were incubated with magnetic streptaividin-coated beads (50uL of slurry)/each RNA probe) for 30’ at room temperature, under agitation (600 RPM in a ThermoMixer, Eppendorf). 200ug of cell lysates (in Pierce IP lysis buffer; #87787, ThermoFisher), derived from fully differentiated CAD or N2a cells, were then incubated with 3’-desthiobiotinilated- RNA/streptavidin beads at 4°C for 1h under agitation (600 RPM). Final elution was performed in 50uL/pull-down. 20uL of each eluate was then analyzed by M/S.

#### Incubation times

30’ @ RT, 600RPM agitation for the binding of the labeled RNA to the beads; 1h @ 4C, 600RPM agitation for the RBPs to the RNA and 15’ @ 37C, 600RPM agitation for the elution.

T7IVT-302-F TAATACGACTCACTATAGGagtggaggttcgcccc
IVT-302-R aaacgagaaggcgtggcc
T7IVT-2034-F TAATACGACTCACTATAGGtatttattcaaatagcgtgagg
IVT-2034-R gcatacacaactattaaaagc
T7IVT-2080-F TAATACGACTCACTATAGGtctagggaattcctggctc
IVT-2080-R ccccacattaataagaactaaaaac
T7IVT-2535-F TAATACGACTCACTATAGGctctactgcacttagactctcc
IVT-2535-R attatcaataatttgtcagctaagg
T7_LocMotif_For TAATACGACTCACTATAGGcctccccccccccctgt
LocMotif_Rev ctgcagcaggcaggggcc

The sequence motifs 302; 2034; 2080 and 2535 (Table S3) served as negative controls and were used in equimolar combination (1:1:1:1).

### Mass spectrometry

#### The pulldown samples were subjected to Trichloroacetic (TCA) precipitation

20 μl of each sample + 80 μl of H2O + 100 μl of 10% TCA (5% TCA end concentration). The resulting protein pellets were washed twice with cold acetone, dried and dissolved as follows: 45 μl of 10 mM Tris/2 mM CaCl2, pH 8.2 buffer; 5 μl trypsin (100 ng/μl in 10 mM HCl); 0.3 μl trypsin Tris 1M, pH 8.2 to adjusted to pH 8. The samples were then processed with microwave-assisted digestion (60° C, 30 min) and dried. The dried digested samples were dissolved in 20 μl ddH2O + 0.1% formic acid; transferred to the autosampler vials for Liquid chromatography-mass spectrometry analysis (LC-MS/MS);

2 μl were injected on a nanoAcquity UPLC coupled to a Q-Exactive mass spectrometer (Thermo Scientific).

The protein identification and quantification was performed using MaxQuant v1.6.2.3 and the data were searched against the Swissprot mouse database. The mass spectrometry proteomics data have been deposited to the ProteomeXchange Consortium via the PRIDE partner repository (Perez-Riverol et al., 2019) with the dataset identifier PXD025492.

### Single molecule fluorescence in situ hybridization

smFISH staining was performed according to a previously published protocol (Borrelli and Moor, 2020) with minor adaptations. Briefly, 50’000 CAD or N2A cells were seeded per well in 24-well plate and grown on poly-D- lysine (Gibco, A3890401) coated coverslips (Thermo Scientific A67761333) with cell differentiation medium (Day 1). One day after transfection (Day 5), cells were flushed with cold PBS and fixed in 4% paraformaldehyde (PFA, Santa Cruz Biotechnology, sc-281692) in PBS for 10 min and subsequently washed two times with cold PBS. Fixed cells were permeabilized with 70% ethanol for at least 1 hour or maintained overnight at 4 degree (Day 6). The permeabilized samples were washed once with wash buffer A (10% Formamide (Ambion, 9342), 20% Stellaris RNA FISH Wash Buffer A (Biosearch Technologies Cat# SMF-WA1-60) in nuclease-free water (Ambion, AM9932)) for 5 min each at 37 °C. Probe libraries were designed using the Stellaris FISH Probe Designer (Biosearch Technologies, Inc., Petaluma, CA, see Table S7) and covalently coupled to Cy5 (Lyubimova et al., 2013).

Hybridization mix (200 nM probes in Stellaris hybridization buffer, Biosearch Technologies Cat# SMF-HB1-10 and 10% Formamide) was added after aspirating the wash buffer A and the sample was incubated upside-down facing the buffer at 37°C in the dark for about 18–24 h (within assembled humidified chamber). Hybridization mix was carefully removed and sample was washed once with wash buffer A at 37 °C for 30 min each in the dark. Samples were stained with DAPI (Invitrogen, D1306; 10 μg/ml in wash buffer) for 30 min at 37 °C in the dark. DAPI solution was aspirated and samples were washed once with wash buffer B (Biosearch Technologies Cat# SMF-WB1-20) for 5 min. Samples on cover glass were gently mounted upside-down with a small drop of ProLong^™^ Gold (Invitrogen^™^ P36930). smFISH imaging was performed on a Leica THUNDER Imager 3D Cell Imaging system. 100x NA=1.4 oil immersion objective lens was used. smFISH-based quantifications of neurite localization were carried out as follows: We measured fluorescence intensity of the smFISH signal along the dendrite, starting from the cell body. We then applied background subtraction and normalized the values for expression level (cell-body fluorescence intensity). A measure for neurite localization was obtained by taking the fold change in mean signal in distal parts of a neurite (more than 30 μm from the cell body) over the mean signal in proximal parts of the same neurite (between 5 and 30 μm from the cell body).

smFISH-based quantification of nuclear/cytoplasmic ratio was carried out by manually segmenting nuclear, soma cytoplasmic, and intercellular regions based on the DAPI and GFP background channels in Fiji (Schindelin et al., 2012). The intercellular background signal intensity in the Cy5 channel was then subtracted from both the nuclear and cytoplasmic regions of interest before calculating the nuclear / cytoplasmic ratio for each segmented cell.

### Mapping next generation sequencing reads and computing enrichment scores

Mapping was performed using custom-made Python scripts. To unambiguously identify the library variant, a unique 12-mer barcode sequence was placed at the 5’ end of each variable region. DNA was sequenced on a NovaSeq 6000 sequencer (SP flow cell, paired end: read 1 30 bp, read 2 84 bp) and demultiplexed using bcl2fastq. We used read 2 to determine for each read its variant barcode and discarded all the reads that could not be assigned to a library variant of origin. Furthermore, we extracted the corresponding UMI from read 1 and used the UMI count per library variant per sample as the starting point for all subsequent analyses.

Enrichment (logFC(neurite/soma) and associated p-value) was calculated using edgeR (Robinson et al., 2010) (version 3.28.1; based on fitting a generalized linear model and performing likelihood ratio test to test for enrichment) based on UMI counts of each library sequence in the neurite and soma fraction of three biological replicates.

### RBP motif enrichment analyses

RBP motif enrichment analysis In order to find enriched RBP motifs in published localized RNAs, we downloaded 103 position frequency matrices (PFMs) that correspond to 85 human RBPs from the RNAcompete paper (Ray et al., 2017). These PFMs (which are of length seven or eight) are generated from the alignment of top 10 7-mers determined using all data (i.e. both setA and setB of RNAcompete pool). Rather than using these top 10 7-mers directly, we generated the top 10 n-mers from the PFMs. In this way, we were able to scan for motifs that are longer than seven. An example is the FXR1 RBP for which the PFM inferred by RNAcompete is of length eight. By using the top 10 8-mers in our motif search, we can represent the binding preferences to all eight positions of this PFM. Next, we have collected a set of dendritically localized RNAs and background RNAs from Middleton et al (Middleton et al., 2019), as well as a second set of RNAs localized in Neurite versus Soma from Zappulo et al (Zappulo et al., 2017) (logFC > 1 for RNAs localized in Neurite and logFC> -2 for RNAs localized in Soma to have a comparable set of RNAs in terms of numbers of RNAs in each set). 3’UTRs of the reported localized RNAs were extracted from Ensemble using biomaRt R package (Durinck et al., 2009) for the RNA ids with corresponding match in the database. All extracted 3’UTR sequences were converted to “BStringSet” using Biostring R package (Pages et al., 2014) for the downstream analysis. To calculate the enrichment of each RBP motifs in RNAs reported as localized in Dendrites/Neurites versus Background/Soma in Middleton et al. and Zappulo et al correspondingly, we used enrich_motifs function from universalmotif R package (Tremblay) which provided the number of motif hits for each of the RBP top 10 n-mers as well as the corresponding p-value and q-value.

### De-novo motif analysis

In addition to the enrichment of known RBP motifs we scanned published localized RNA 3’UTRs for de-novo motifs. To do so, for each set of extracted 3’UTRs from localized RNA in Middleton et al. and Zappulo et al. papers we used MEME Suite for the following analysis: i)MEME de-novo motif discovery analysis (Bailey, 2003) to discover novel, ungapped motifs in our localized sets of RNAs. We ran this function in a Differential Enrichment mode by providing the Dendrite/Neurite localized 3’UTRs as primary sequences and Background/Soma RNAs as control sequences. We chose anr (Any Number of Repetition) as site distribution of the function and we looked for the 20 de-novo motifs. The rest of the parameters were kept as default. ii) MAST Motif scanning analysis (Bailey and Gribskov, 2000) on the set of motifs we had discovered de-novo from the previous step. We provide the MEME xml output as the motif set for the MAST function and the 3’UTR of the Dendrite/Neurite localized RNAs as the sequence sets to scan for the matches to motifs.

### Prediction of localization

Machine learning procedures were carried out using the python scikit-learn and XGBoost package. Initially, from all duplicated sequences (e.g. barcode control sets), which passed filtering, a single variant was randomly chosen for all subsequent steps to avoid biases resulting from having duplicated sequences. 10% of the variants were put aside and used only for evaluation of models built using the other 90% (80%). We chose Gradient Boosting Decision Trees (XGBoost, (Chen and Guestrin, 2016)) as the prediction algorithm because it can capture non-linear interactions between features and has proven to be a powerful approach in predicting the effect of regulatory regions (Mikl et al., 2019, 2020).

We used two sets of features for our prediction: 1) We computed counts of all possible 4mers in each library sequence (not taking into account the barcode and constant sequences like primer binding sites). 2) We computed cumulative binding scores of a set of 218 RBPs (RNAcompete, (Ray et al., 2017)) for each library sequence (not taking into account the barcode and constant sequences like primer binding sites).

#### We trained two models

one predicting neurite enrichment, where the positive class was defined as having a logFC>0 and an associated p-value<0.05 in both CAD and Neuro-2a cells. The second model was trained to predict soma enrichment, where the positive class was defined as having a logFC<0 and an associated p-value<0.05 in both CAD and Neuro-2a cells. For prediction of the localization behavior of unseen variants, we used the difference in the predicted probability for the positive class between the two models (P(is neurite enriched) - P(is soma enriched)) as the final output.

For prediction of the localization behavior of native transcripts we predicted the probability of neurite localization (as described for the test set above) for all potential 3’UTR tiles of a gene with length 150 bp and a step size between tile starting points of 50 bp (i.e. tile 1 corresponds to 3’UTR positions 1-150, tile 2 to 3’UTR positions 51-200, etc.). To capture the positive contributions to neurite localization, we calculated from all individual tiles of a 3’UTR the median prediction probability of the model predicting neurite localization. From this we subtracted the maximal prediction probability of the model predicting soma localization, accounting for the fact that according to our results even single soma-restriction signals can overrule other signals promoting neurite localization. We then compared this combined model output to the localization behavior (logFC>0 or <0) reported by Taliaferro et al. (2016).

### General data analysis

For data analysis, we used python 3.7.3 with pandas 0.24.2, numpy 1.16.2, seaborn 0.9.0, scipy 1.2.1, scikit-learn 0.20.3 and shap 0.34.

## Data availability

Illumina sequencing data generated in this study are available in the NCBI gene expression omnibus (GEO) under accession GSE173098. The mass spectrometry proteomics data have been deposited to the ProteomeXchange Consortium via the PRIDE partner repository with the dataset identifier PXD025492.

## Author contributions

Conceptualization and Methodology: M.M. and A.E.M..; Software: M.M., A.L. and S.B.; Formal Analysis: M.M. and A.L.; Investigation: M.M., D.E., M.L., F.M. and K.H.; Funding Acquisition: A.E.M.; Writing, Visualization and Supervision: M.M. and A.E.M.

## Notes

### Competing Interest Statement

The authors have declared no competing interest.

