## Supplementary figures and images for "A massively parallel reporter assay reveals focused and broadly encoded RNA localization signals in neurons"

### SupplementaryFigures_RNAlocMPRA.pdf

Figure S1

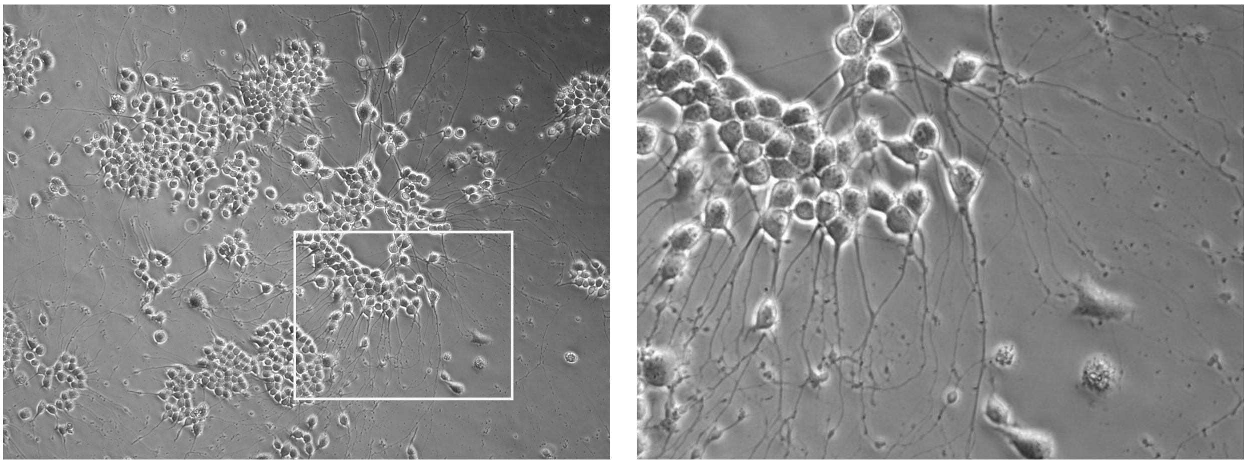

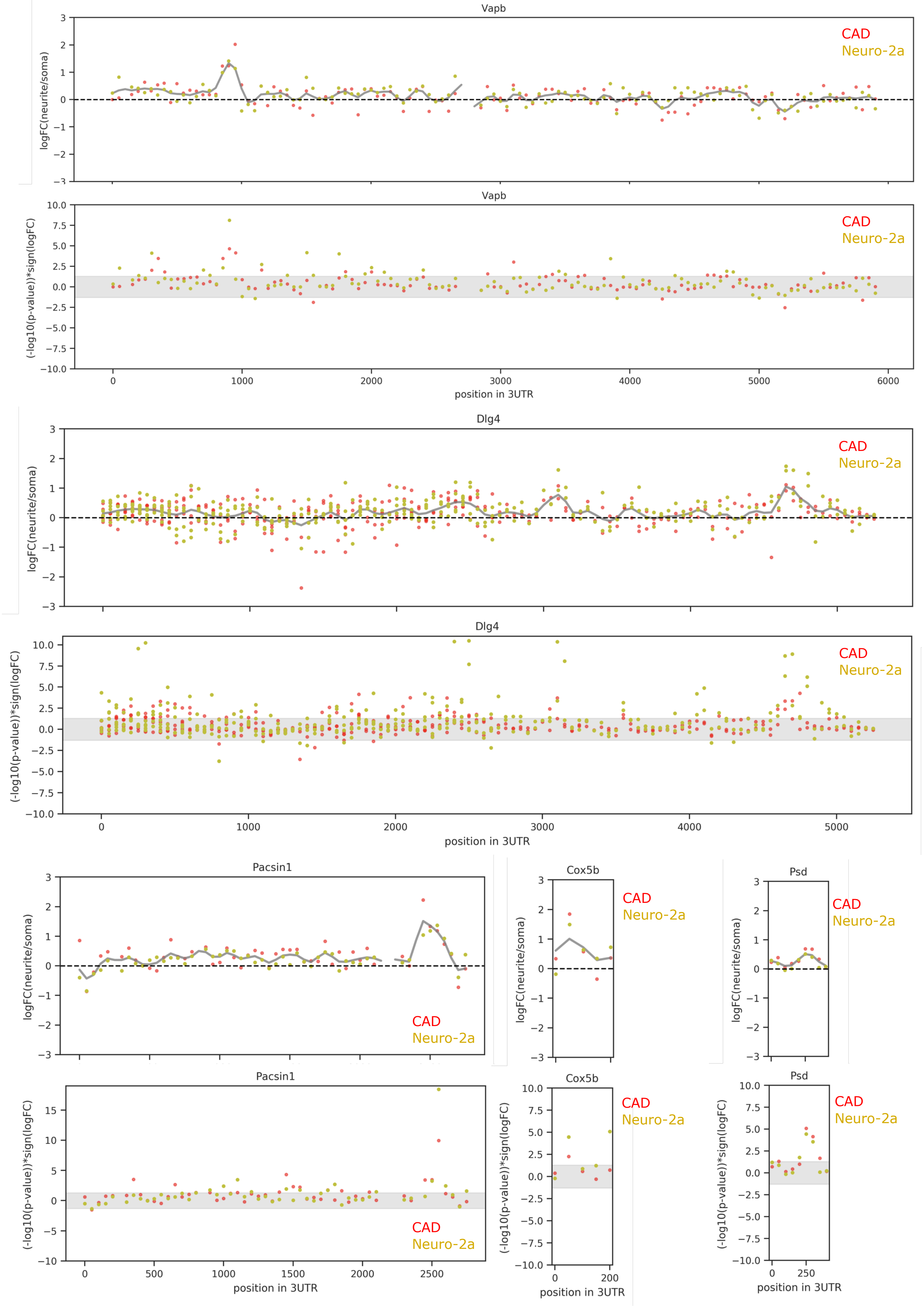

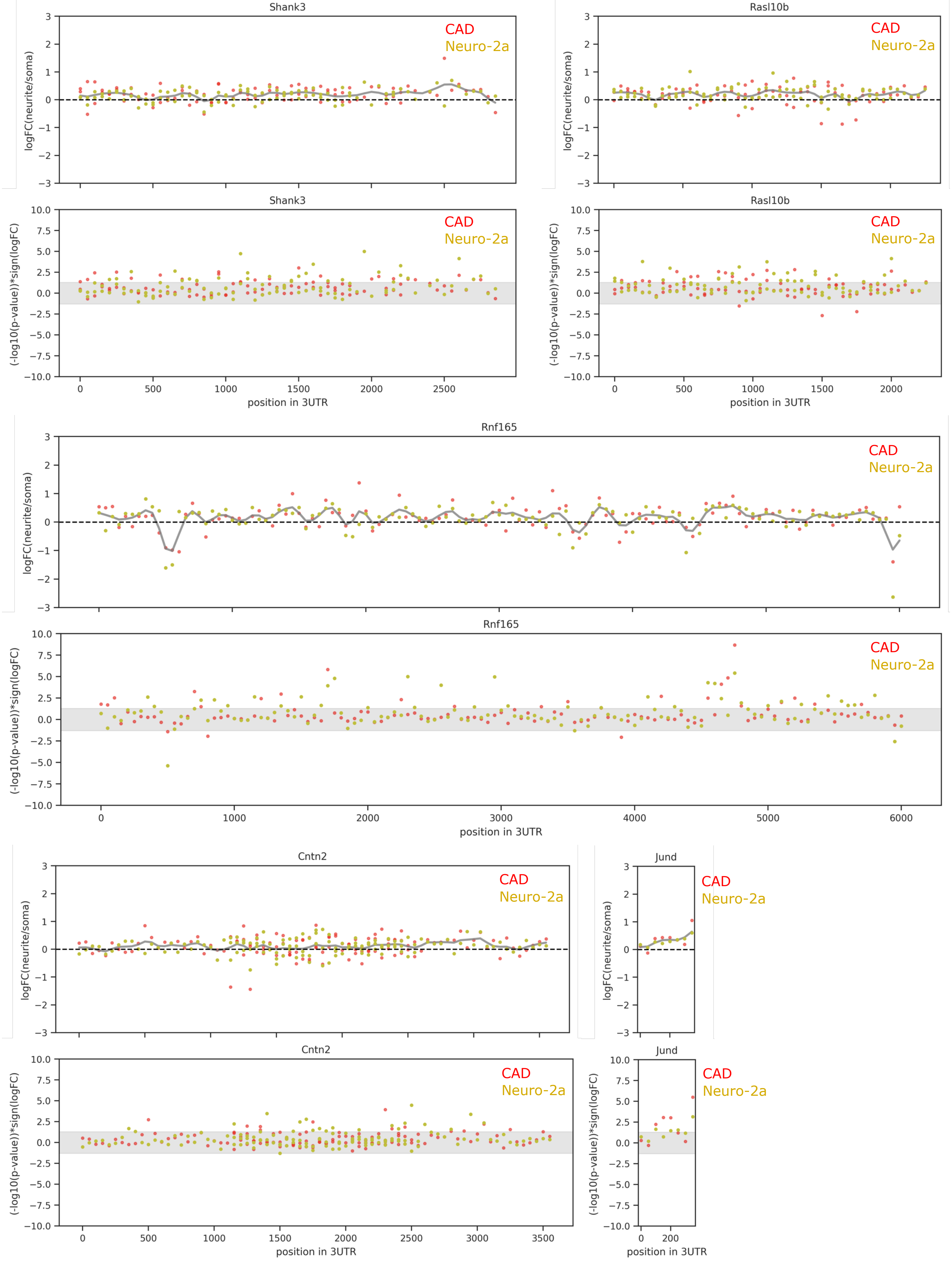

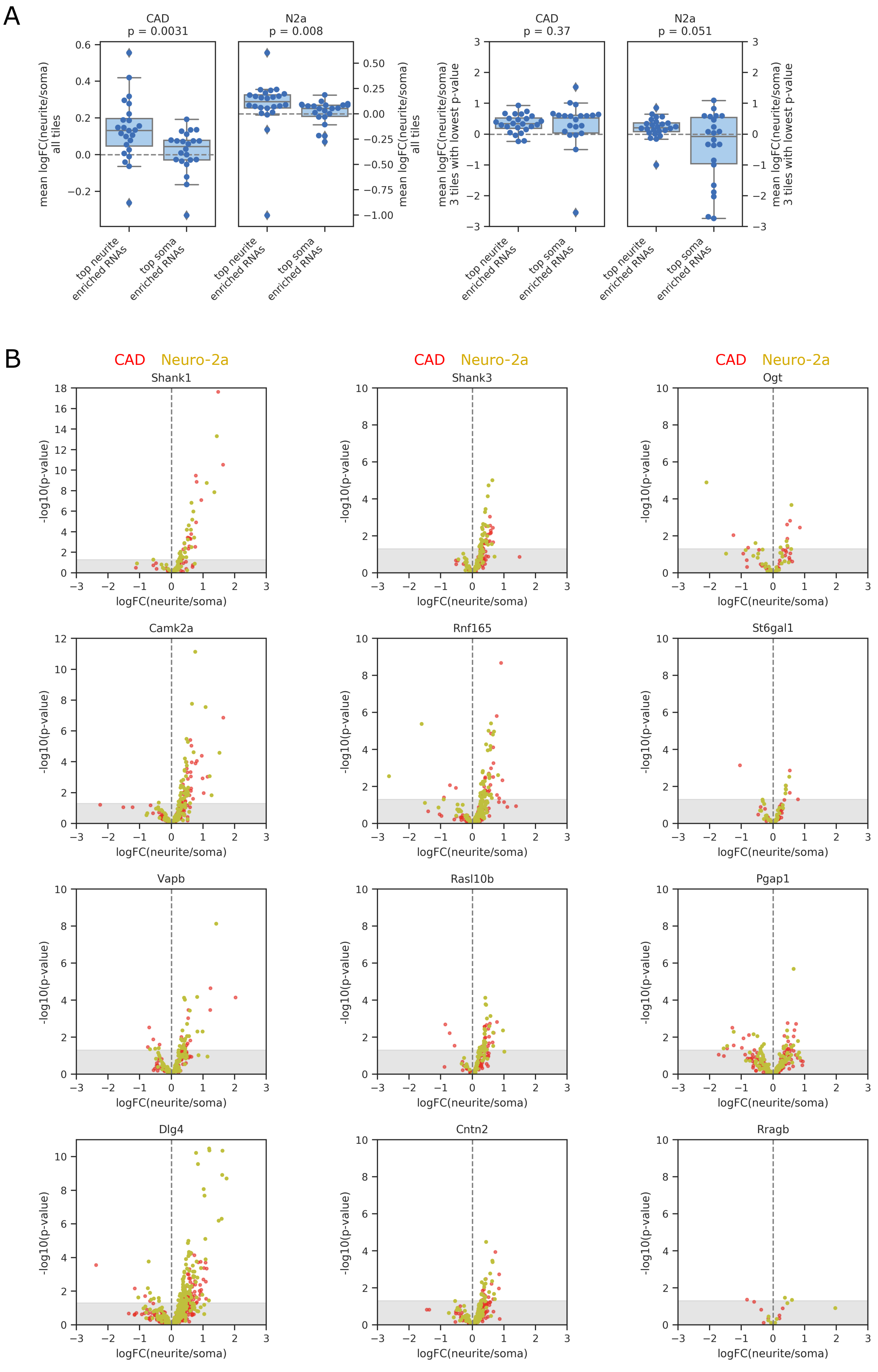

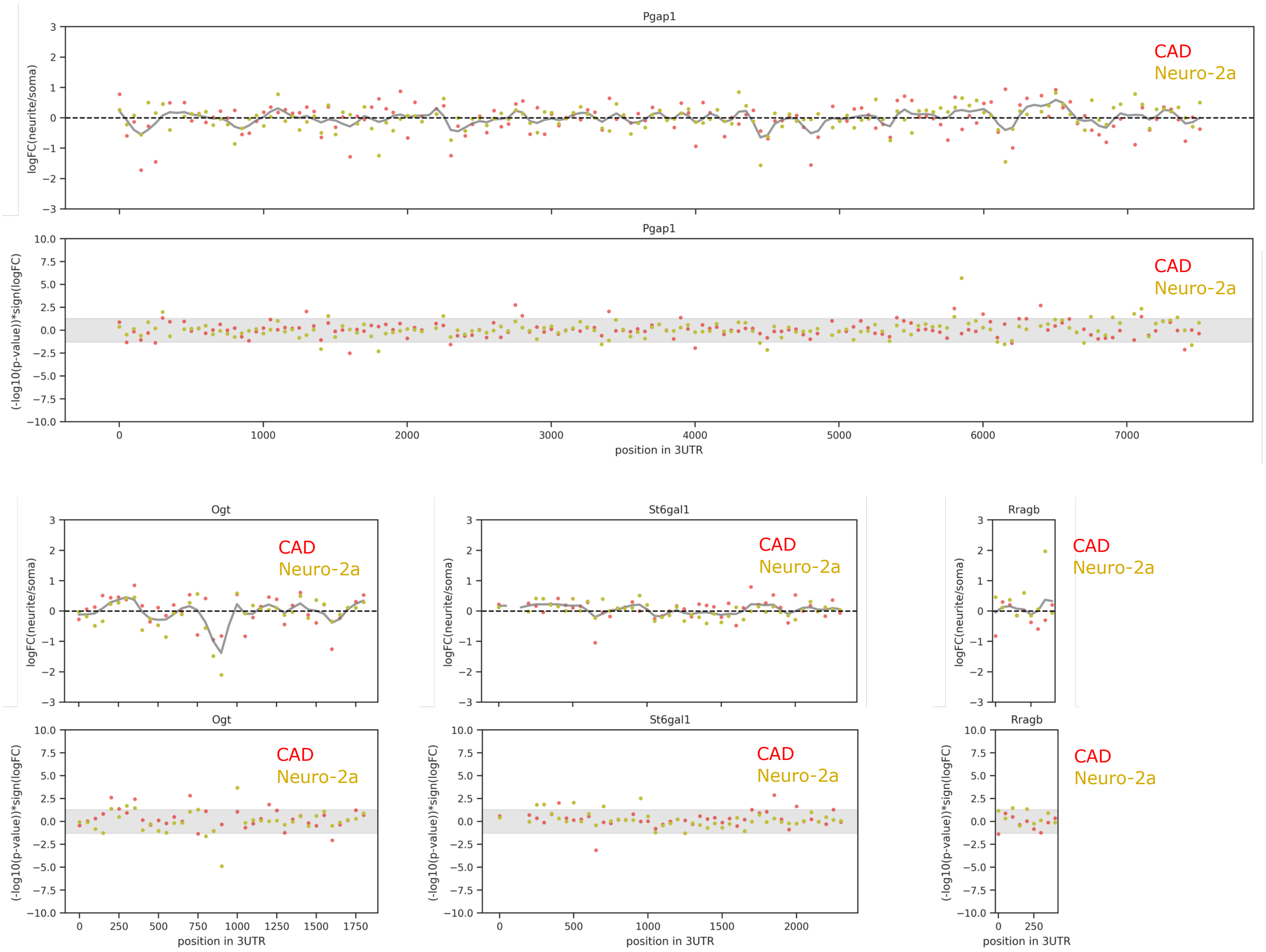

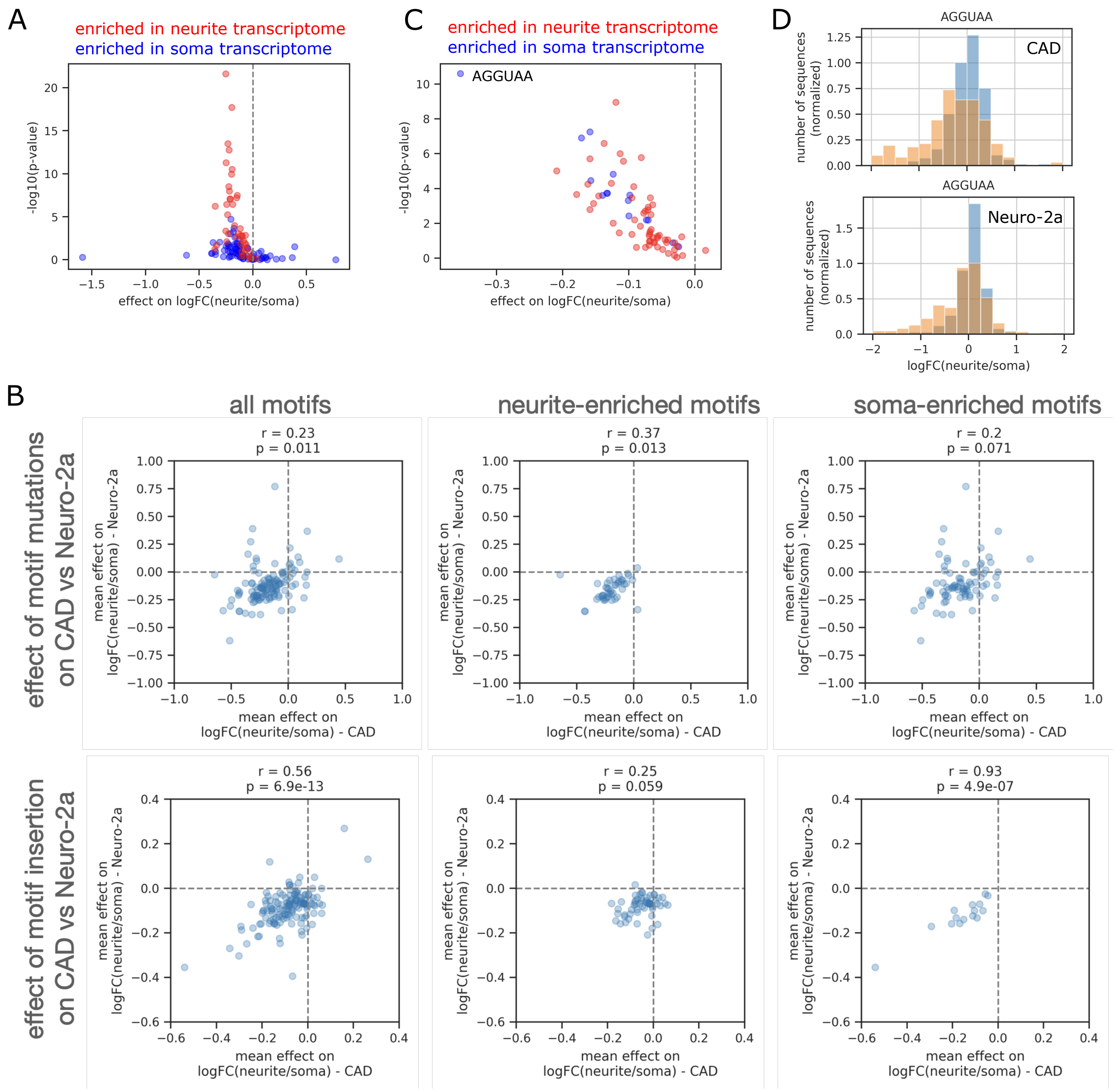

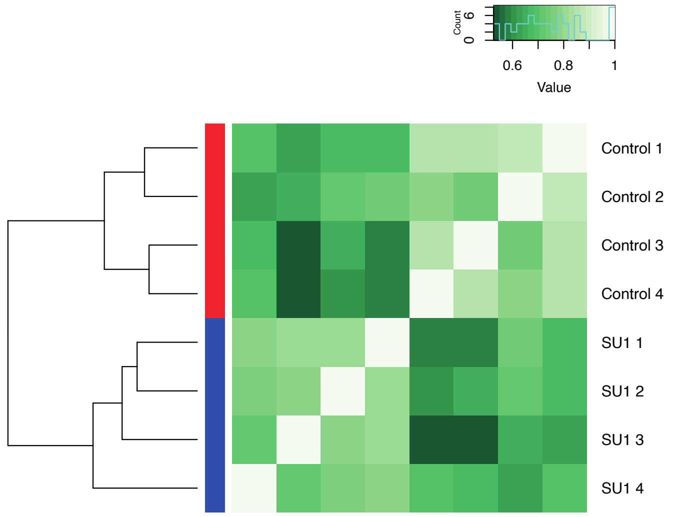

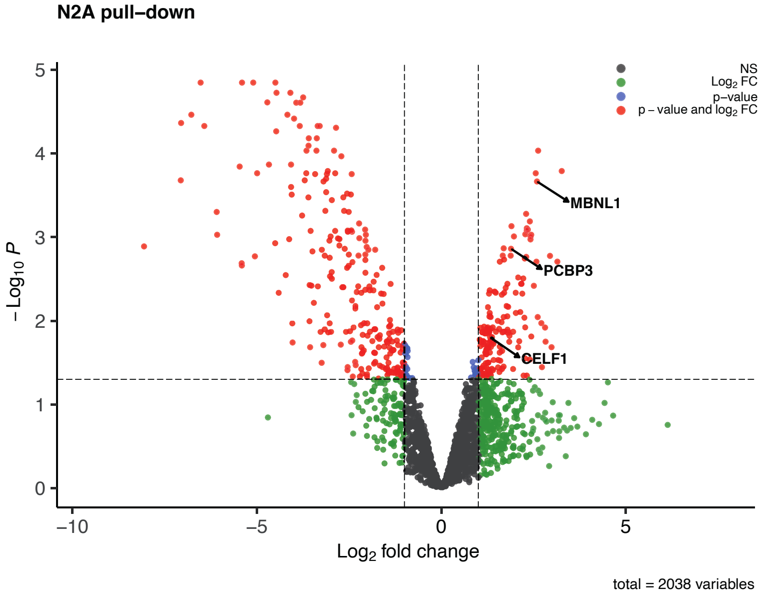

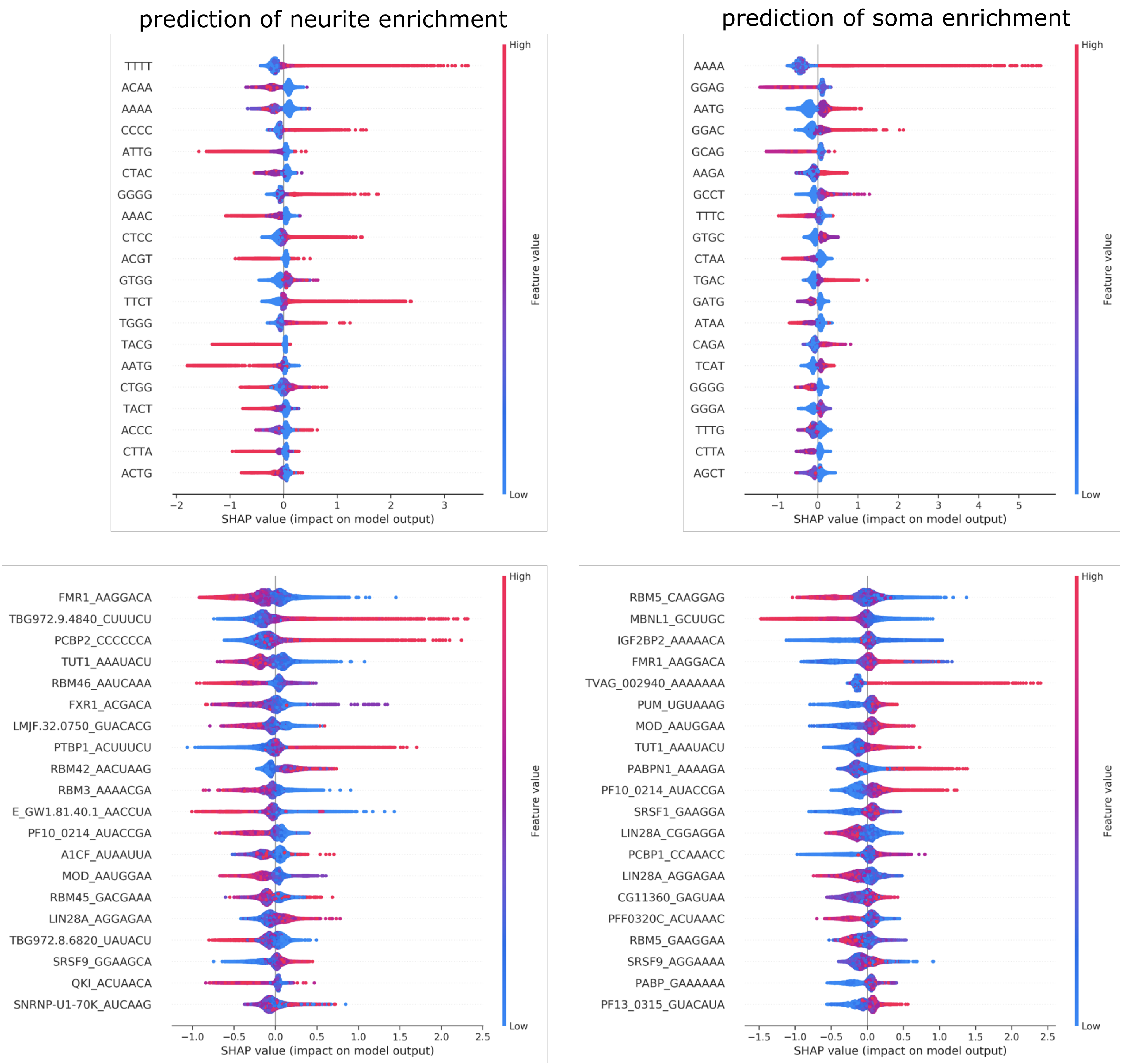
